# Lysosomal RNA profiling reveals targeting of specific types of RNAs for degradation

**DOI:** 10.1101/2025.09.09.674968

**Authors:** G. Jordan Ray, Eleonora Nardini, Heather R. Keys, Daniel H. Lin, David M. Sabatini, David P. Bartel

## Abstract

Autophagy targets a wide variety of substrates for degradation within lysosomes^1^. While lysosomes are known to possess RNase activity^2^, the role of lysosomal RNA degradation in post-transcriptional gene regulation is not well understood. Here, we define RNASET2, PLD3, and both endogenous and exogenous RNase A family members as lysosomal RNases. Cells lacking these RNases accumulated large amounts of lysosomal RNA. Although all types of RNA can be found within lysosomes, SRP RNAs, Y RNAs, 5′ TOP mRNAs, long-lived mRNAs, and mRNAs encoding membrane and secreted proteins were specifically enriched. All types of RNA depend on autophagy for lysosomal targeting, but the lysosomally-enriched RNAs are more sensitive to loss of autophagy, implying that selective mechanisms mediate their lysosomal entry. RNA stability measurements revealed that lysosomally-degraded transcripts also had autophagy-dependent changes in stability. In exploring how specific RNAs are targeted for lysosomal degradation, we found that the Alu domain of SRP RNAs is sufficient for targeting these RNAs to lysosomes in fashion that depends on its interactions with the SRP9 and SRP14 proteins. For mRNAs, 5′ TOP motifs are sufficient to increase their targeting to lysosomes for degradation in a LARP1-dependent manner. Altogether, our results establish lysosomes as selective modulators of cellular RNA content.

## Main Text

Lysosomes are degradative organelles that contain a broad range of hydrolase activities that can break down diverse substrates into molecular building blocks to be recycled^3^. One of the hallmark enzymatic activities of lysosomes is a ribonuclease (RNase) activity, which has been largely attributed to the acid endonuclease Ribonuclease T2 (RNASET2)^2,4,5^. In mammalian cells starved of nutrients, RNA turnover increases in a lysosome-dependent manner, but the identities of such lysosomally degraded RNAs are not known^6^.

Substrates targeted for lysosomal degradation have been reported to enter lysosomes via various pathways, including macroautophagy, microautophagy, and chaperone-mediated autophagy (CMA). Macroautophagy, hereafter called autophagy, initiates in response to stresses, such as starvation, and entails the encapsulation of substrates within a double-membrane vesicle, the autophagosome, which then fuses with lysosomes to enable substrate degradation^7^. Autophagy can target specific substrates for degradation, and the selectivity of this process has been demonstrated for certain organelles, proteins, and bacteria^1^. In microautophagy, the lysosomal membrane invaginates to internalize cytosolic substrates for degradation^8^. In one type of microautophagy, RNautophagy, the membrane proteins LAMP2C (lysosome-associated membrane protein-2 C) and SIDT2 (a mammalian ortholog of the *C. elegans* SID-1 dsRNA transporter) mediate the lysosomal uptake of RNA^9–13^. Lastly, in CMA the chaperone protein HSC70 binds to a specific motif within substrate proteins and together with LAMP2A (lysosome-associated membrane protein-2 A) initiates translocation of such proteins into the lysosome^14^. The extent to which each of these distinct pathways contributes to lysosomal RNA degradation is currently unknown.

Several studies have recently investigated RNA degradation in the vacuole, an organelle present in organisms such as fungi and plants, which is involved in osmoregulation, the degradation of macromolecules, and the storage of nutrients^15^. In *Saccharomyces cerevisiae*, Rny1, the RNASET2 ortholog, is responsible for the vacuolar degradation of both ribosomal RNAs (rRNAs) and specific mRNAs in an autophagy-dependent manner^16,17^. The authors propose that ribosome-bound mRNAs are targeted for lysosomal degradation, which accounts for the degradation of specific mRNAs, as well as the preferential degradation of rRNA. In *Arabidopsis thaliana*, distinct RNAs have been sequenced from vacuolar preparations, but because this sequencing is not quantitative, whether their abundances increase upon loss of RNS2, the RNASET2 ortholog, or whether autophagy contributes to their vacuolar entry remains unclear^18^.

A variety of mechanisms have been identified by which RNAs are regulated and ultimately decayed^19^. Following their transcription, RNAs can undergo nuclear processing and surveillance, such that defective transcripts are degraded. Then, in the cytoplasm additional quality-control and regulatory mechanisms impact RNA stability. These mechanisms are best characterized for mRNAs, where deadenylation of the poly(A)-tail, which triggers decapping and exonucleolytic decay from the 5′ and/or 3′ ends, largely dictates mRNA stability^20,21^. The mechanisms underlying the cytoplasmic degradation of other RNAs is less well studied.

In this study, we investigated lysosome-mediated degradation of endogenous RNAs in mammalian cells, elucidated mechanisms that account for the selective lysosomal degradation of particular RNAs, and established lysosomal RNA degradation as a mechanism for regulating RNA stability.

### RNAs accumulate in lysosomes following RNASET2 loss and mTORC1 inhibition

To identify RNAs degraded within lysosomes, we modified the LysoIP method for isolating lysosomes, which was originally developed for their metabolomic and proteomic analyses^22,23^. In this method, magnetic beads coupled to an anti-HA antibody are used to isolate intact lysosomes from dounce-homogenized lysates of cells expressing a lysosome-specific protein, TMEM192, tagged with three HA epitopes. To eliminate RNAs on the outside of lysosomes, we added an RNase-digestion step immediately after the immunoprecipitation (IP) (Fig. 1a and Extended Data Fig. 1a). As the identities of RNAs inside and outside lysosomes are not known, we first tested the RNase approach on mitochondria isolated by the analogous MitoIP method from human embryonic kidney (HEK) 293T cells^24^. Gratifyingly, in MitoIP samples RNase treatment eliminated cytoplasmic but not mitochondrial rRNAs, indicating that with our method luminal RNAs remain protected while those on the outer membrane are degraded (Extended Data Fig. 1a–c).

**Fig. 1:**
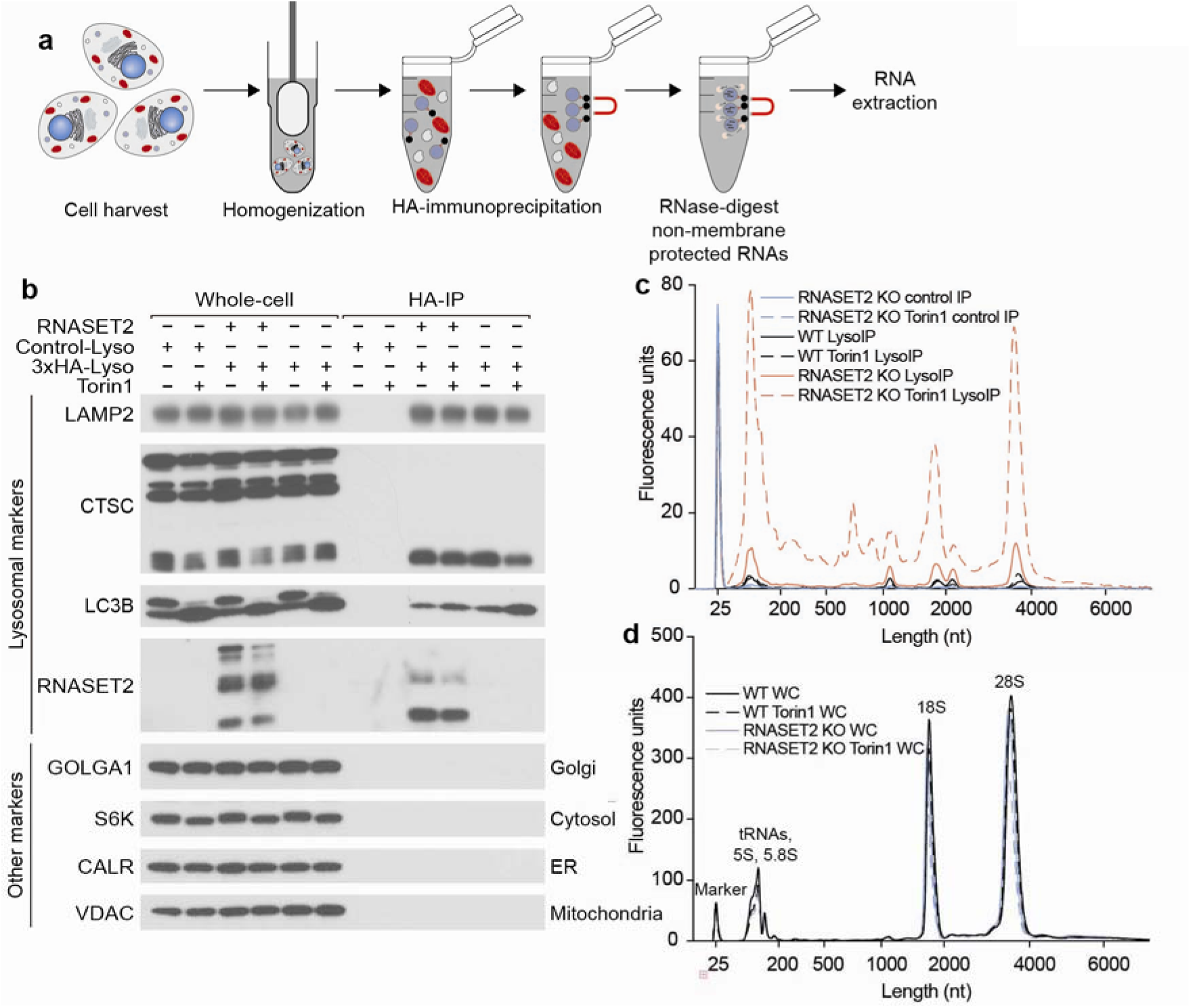
RNAs accumulate in lysosomes following RNASET2 loss and mTORC1 inhibition. **a,** Schematic of lysosomal RNA purification. HEK293T cells were dounce-homogenized to disrupt their plasma membrane while keeping organelles intact, followed by an HA-immunoprecipitation to isolate lysosomes, an RNase I treatment to remove RNAs on the outside membrane, and RNA extraction with TRIzol LS. **b**, Lysosomal purification efficiencies across the indicated genotypes and treatments. Lysosomal capture efficiency and purity were assessed by immunoblotting for protein markers of various subcellular compartments in whole-cell lysates, control immunoprecipitates, or purified lysosomes. Equal numbers of HEK293T cells stably expressing TMEM192-2x-FLAG (Control-Lyso) or TMEM192-3x-HA (3xHA-Lyso) were subjected to the LysoIP protocol. Where indicated, cells were treated with 250 nM Torin1 for 5 h. **c,** Effect of RNASET2 KO and Torin1 on lysosomal RNA accumulation. Lysosomal RNAs from equal proportions of immunoprecipitates from (**b**) were purified and resolved on a bioanalyzer. The peak at 25 nt is a marker. Where indicated, cells were treated with 250 nM Torin1 for 5 h. **d**, Analyses of whole-cell RNA purifications. Equal proportions of whole-cell RNA samples from the same cells used in (**b**) were resolved on a bioanalyzer. The peak at 25 nt is a marker. Other well-established peaks are tRNAs and rRNAs, as marked above each peak. Where indicated, cells were treated with 250 nM Torin1 for 5 h.

Using this modified LysoIP method, we investigated the lysosomal accumulation of RNA in HEK293T cells. We reasoned that the abundance of lysosomal RNA could be increased by lowering lysosomal RNA degradation and/or increasing the flux of RNA into lysosomes. To reduce lysosomal RNase activity, we generated RNASET2 knockout (KO) HEK293T cells, and to increase flux into the lysosome, we activated autophagy by inhibiting mTOR Complex 1 (mTORC1) with Torin1^25^. Importantly, the modified LysoIP method captured equivalent amounts of lysosomes across the genotypes and treatments, as judged by the lysosomal membrane and luminal markers LAMP2 and Cathepsin C (CTSC), respectively (Fig. 1b). In wild-type (WT) cells, the LysoIP captured more RNA than a mock IP, but mTORC1 inhibition only minimally affected the amount of lysosomal RNA recovered (Fig. 1c). This observation suggests that in WT cells, the rate of RNA degradation within lysosomes is faster than the flux of RNA into lysosomes, even upon activation of autophagy. In contrast, in RNASET2-KO cells, basal lysosomal RNA levels substantially increased, and even more lysosomal RNA accumulated after mTORC1 inhibition (Fig. 1c). These results show that RNA is constitutively degraded within lysosomes and that the process can be augmented by reduced mTORC1 activity.

To ask if most RNAs enter lysosomes via processes that depend on vesicle fusion, we used the vacuolar sorting protein 34 (VPS34) inhibitor SAR405, which inhibits the fusion of autophagosomes with lysosomes^26^. When combined with Torin1, SAR405 negated the effect of mTORC1 inhibition, indicating that vesicle fusion and autophagy are necessary for targeting RNAs for lysosomal degradation (Extended Data Fig. 1e–f and 1h).

Lysosomal RNAs in RNASET2-KO cells were generally smaller and of more varied sizes than whole-cell (WC) RNA (Fig. 1c–d), which could be a result of specific RNAs being targeted for lysosomal degradation, degradation of RNA still happening within RNASET2-KO lysosomes, or both. To determine if RNA degradation still occurs in lysosomes lacking RNASET2, we treated RNASET2-KO cells with Torin1 for 2 h to initiate the trafficking of RNAs to lysosomes and then with SAR405 to prevent further vesicle fusion. If minimal lysosomal RNase activity remained in RNASET2-KO cells, then lysosomal RNA levels would be expected to remain constant after SAR405 treatment. However, this was not the case; lysosomal RNA levels declined post addition of SAR405, consistent with the presence of other lysosomal RNases (Extended Data Fig. 1g–h).

### A CRISPR screen identifies PLD3 as a lysosomal RNase

Given the ability of RNASET2-KO cells to still degrade lysosomal RNA, we sought to identify other factors responsible for lysosomal RNA degradation. To this end, we established a fluorescence-activated cell sorting assay compatible with CRISPR loss-of-function screening. This assay was inspired by the observation that a Zebrafish mutant for RNASET2 exhibits increased staining by the dye acridine orange, which is often used to quantify acidic organelles^27,28^. We validated that RNASET2-KO HEK293T cells treated with Torin1 had an increased acridine orange signal (ratio of red to green) compared with KO cells in which we restored expression of RNASET2 (Fig. 2a).

**Fig. 2:**
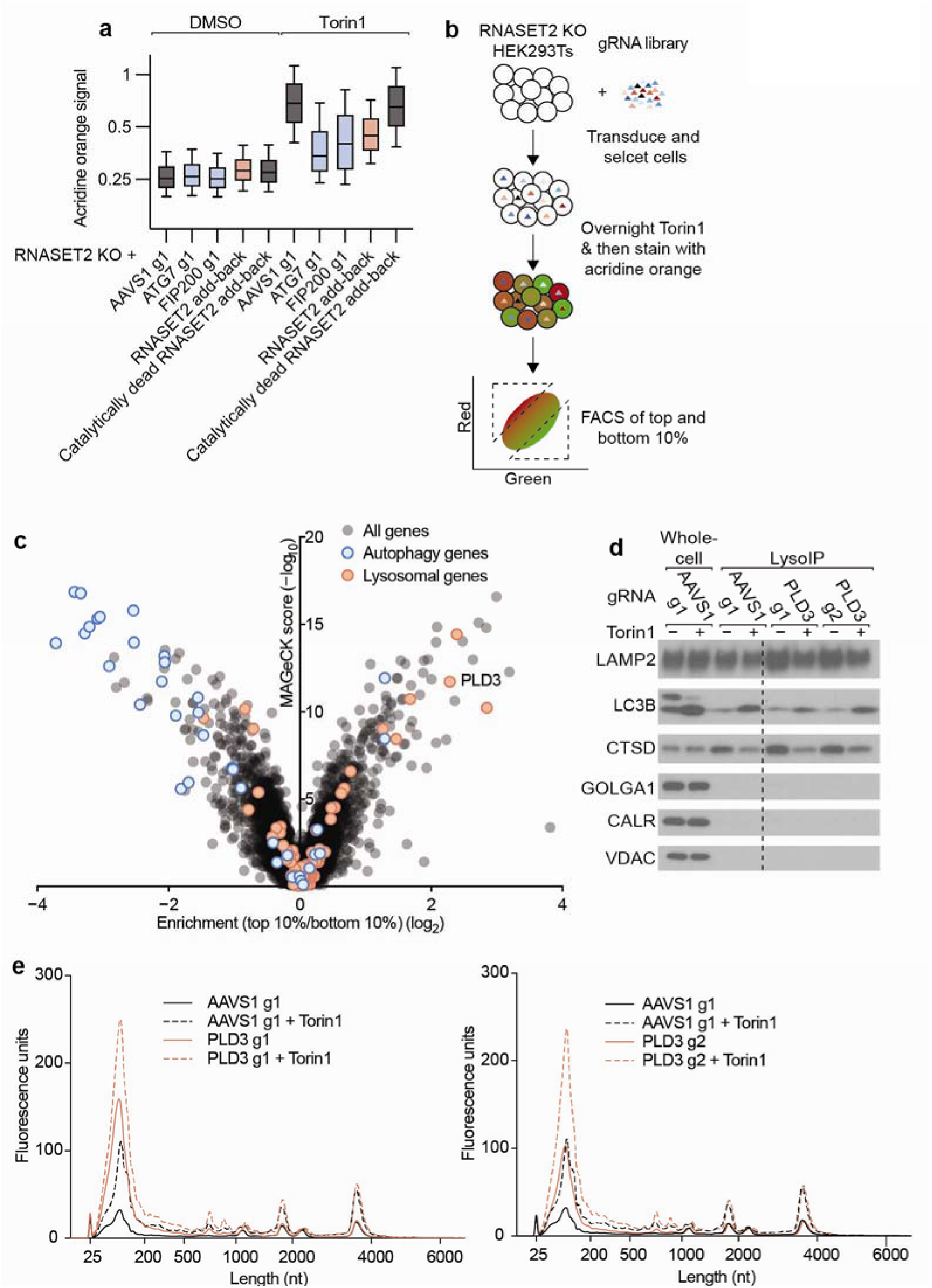
A CRISPR screen identifies PLD3 as a lysosomal RNase. **a**, Acridine orange FACS assay. RNASET2-KO cell lines stably expressed either gRNAs and Cas9, or RNASET2 rescue constructs with or without catalytically dead H65F/H118F mutations. Cells were stained with 4 ug/ml acridine orange and subjected to flow cytometry to measure red and green fluorescence. Acridine orange signal is the ratio of red-to-green fluorescence. Where indicated, cells were treated with 250 nM Torin1 overnight. **b,** Schematic of the acridine orange screen. RNASET2-KO cells were infected with a genome-wide CRIPSR KO library. At 13 and 17 days after transduction, cells were treated with 250 nM Torin1 overnight, stained with 4 ug/ml acridine orange, and the top and bottom 10% of cells were sorted based on their red-to-green ratio. **c,** Results of the acridine orange screen. Guide-RNA enrichment in cells with red-to-green ratios in the top 10% relative to that in cells with ratios in the bottom 10%, is plotted as a function of MAGeCK score (Supplementary Table 1). Genes encoding lysosomal proteins were defined as those with LysoIP proteomic enrichments higher than 3 from Wyant *et al.*^23^. **d**, Lysosomal purification efficiencies across the indicated genotypes and treatments. Lysosomal capture efficiency and purity were assessed by immunoblotting for protein markers of various subcellular compartments in whole-cell lysates, or purified lysosomes. Equal numbers of RNASET2-KO HEK293T cells stably expressing TMEM192-3x-HA, Cas9, and the indicated guide RNA, were subjected to the LysoIP protocol. Where indicated, cells were treated with 250 nM Torin1 for 5 h. **e**, Effect of PLD3 loss on lysosomal RNA degradation. Lysosomal RNAs from equal proportions of immunoprecipitates from (**d**) were purified and resolved on a bioanalyzer. The peak at 25 nt is a marker. Where indicated, cells were treated with 250 nM Torin1 for 5 h.

Furthermore, this increased acridine orange signal was dependent upon the two autophagy proteins autophagy-related protein 7 (ATG7) and Focal adhesion kinase- interacting protein of 200 kDa (FIP200), consistent with our SAR405 results (Fig. 2a).

We transduced RNASET2-KO HEK293T cells with a genome-wide CRISPR-Cas9 KO library that contained five guides per gene^29^. Then, 13 and 17 days after transduction, cells were treated overnight with Torin1, stained the next day with acridine orange, and sorted to isolate those in the top and bottom ten percentiles of the acridine orange signal (Fig. 2b). Genomically integrated guide RNA sequences were then amplified and sequenced from each of these isolated populations to identify how loss of each gene affected the acridine orange signal. As expected, loss of most autophagy factors led to a decreased acridine orange signal (Fig. 2c and Supplemental Table 1). To identify factors responsible for lysosomal RNA degradation, we cross-referenced the set of proteins previously reported to be preferentially localized to the lysosome^23^ with the proteins that increased acridine orange signal upon depletion in our screen. Among the top three candidates was the lysosomal protein Phospholipase D3 (PLD3) (Fig. 2c and Supplemental Table 1). To validate PLD3 as a lysosomal RNase, we purified lysosomal RNA from RNASET2-KO cells lines expressing Cas9 and guide RNAs targeting PLD3. The lysosomes from these double-KO cells contained both more RNA and altered RNA sizes when compared to lysosomes from RNASET2-KO cells (Fig. 2d–f). These findings were consistent with recent evidence that PLD3 is an endolysosmal DNA and RNA 5′ exonuclease and demonstrated that in our cell system both RNASET2 and PLD3 degrade lysosomal RNA^30–35^.

### Endogenous and exogenous RNase A contribute to lysosomal RNA degradation

We also examined whether exogenous protein factors might contribute to lysosomal RNase activity. For the work presented above, HEK293T cells were cultured in Freestyle media containing 1% serum, but these cells can also grow in serum-free medium, and we noted that in the absence of serum, they contained more lysosomal RNA, which suggested the existence of a serum-derived nuclease activity (Fig. 3a and Extended Data Fig. 2a). To identify factors responsible for this activity, we fractionated serum using cation exchange chromatography and tested fractions for their ability to phenocopy the impact of adding serum to the media on lysosomal RNA accumulation (Fig. 3b and Extended Data Fig. 2b–c). Fractions that recapitulated the effects of serum on lysosomal RNA degradation were enriched for bovine RNase A family members (ANG1, RNASE4, BRN, and ANG2) (Fig. 3c and Supplemental Table 2).

**Fig. 3:**
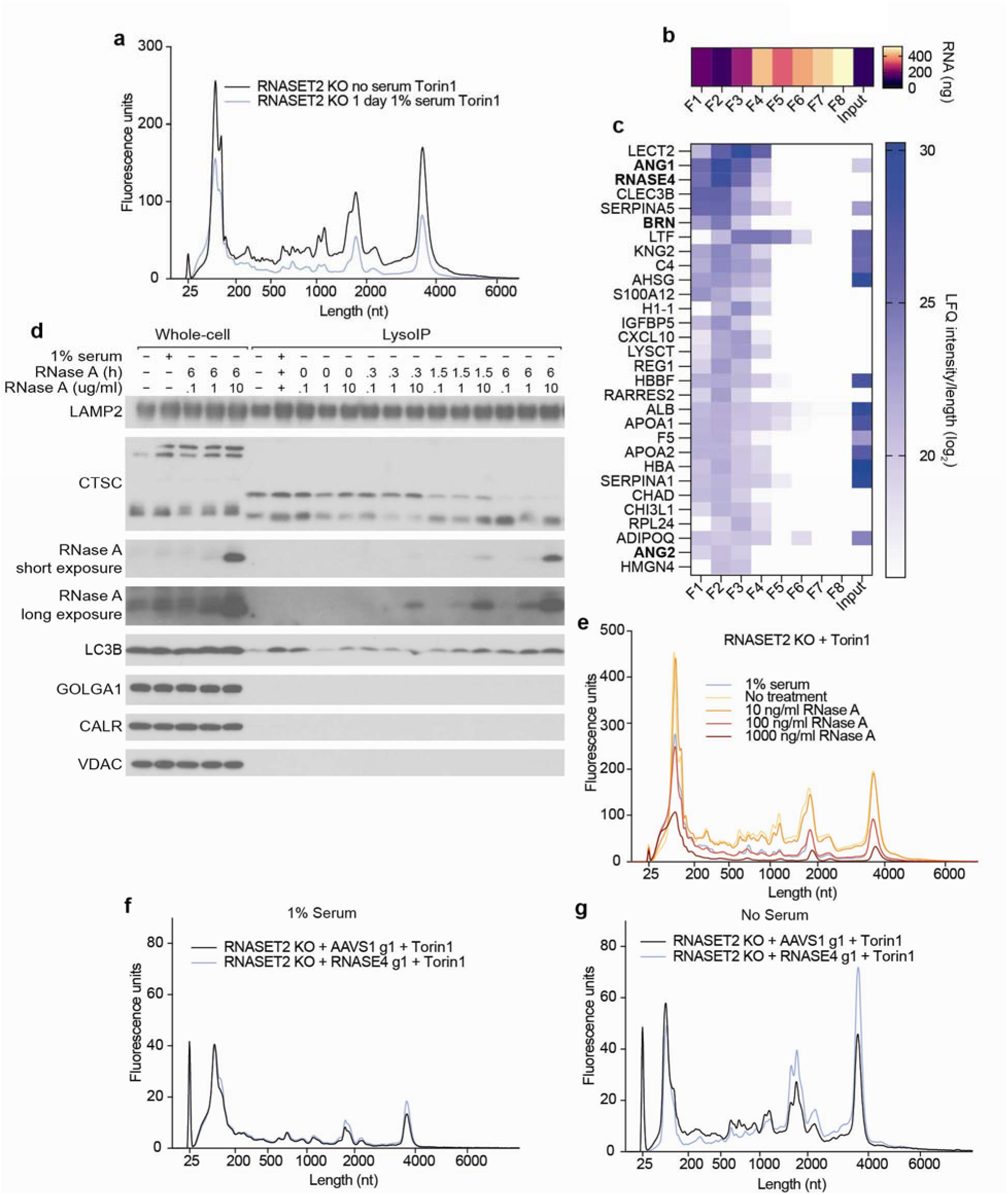
Endogenous and exogenous RNase A contribute to lysosomal RNA degradation. **a**, Effect of media serum on lysosomal RNA accumulation. Lysosomal RNAs from equal proportions of immunoprecipitates from cells that had been cultured with and without serum (Extended Data Fig. 2A) were purified and resolved on a bioanalyzer. The peak at 25 nt is a marker. Cells were treated with 250 nM Torin1 for 6 h. **b,** Effects of fractionated serum (Extended Data Fig. 2B) on lysosomal RNA quantities. Lysosomal RNAs from equal proportions of immunoprecipitates from cells cultured with the indicated serum fractions (Extended Data Fig. 2C) were purified and then quantified on a bioanalyzer. **c**, Proteomics of fractionated serum. Fractionated serum (Extended Data Fig. 2B) was analyzed by proteomics (Supplementary Table 2). Shown are the top 30 bovine proteins ranked by the sum of their length normalized label-free quantification intensity in fractions 1 to 3. RNase A family members are indicated in bold text. **d**, Time- and dose-dependent uptake of RNase A and its lysosomal localization. Lysosomal capture efficiency and purity were assessed by immunoblotting for protein markers of various subcellular compartments in whole-cell lysates, or purified lysosomes. Equal numbers of RNASET2 PLD3 RNASE4 tKO HEK293T cells stably expressing TMEM192-3x-HA were incubated with the indicated concentration of RNase A for the indicated time and then subjected to the LysoIP protocol. **e**, Dose-dependent effect of RNase A on lysosomal RNA degradation. Lysosomal RNAs from equal proportions of immunoprecipitates from cells cultured with the indicated amounts of RNase A (Extended Data Fig. 2D) were purified and resolved on a bioanalyzer. The peak at 25 nt is a marker. RNase A or serum treatment was overnight. Torin1 treatment was 250 nM for 5 h. **f**, Effect of RNASE4 on lysosomal RNA degradation in the presence of serum. Lysosomal RNAs from equal proportions of immunoprecipitates from RNASET2-KO HEK293T cells stably expressing TMEM192-3x-HA, Cas9, and the indicated guide RNA (Extended Data Fig. 2H) were purified and resolved on a bioanalyzer. The peak at 25 nt is a marker. Cells were grown in the presence of 1% serum overnight prior to LysoIP. Torin1 treatment was 250 nM for 6 h. **g,** Effect of RNASE4 on lysosomal RNA degradation in the absence of serum. This panel is as in (**f**) except cells were grown in the absence of serum.

Others have reported RNase A uptake by endocytosis, but the potential physiological function of this phenomenon has not been established^36^. When added to the medium, we found that purified RNase A (bovine RNASE1) was taken up by cells and entered lysosomes in a dose- and time-dependent manner (Fig. 3d). Furthermore, ∼100 ng/mL (7 nM) RNase A phenocopied the effects of adding 1% serum to the medium on lysosomal RNA degradation (Fig. 3e and Extended Data Fig. 2d). As blood from healthy humans contains ∼200 ng/mL RNASE1, this appeared to be a physiologically relevant effect^37^.

To determine if endogenous RNase A homologs also impact lysosomal RNA degradation, we created RNASET2-KO cells also deficient for RNASE4 and ANG (RNASE5), which encode the two secretory RNases expressed in HEK293Ts. When these cells were cultured in the presence of serum, lysosomal RNA degradation was the same as in cells lacking only RNASET2, but when cultured in the absence of serum, lysosomal RNA degradation was reduced (Extended Data Fig. 2e–g). RNASE4 alone accounted for this effect, as its loss phenocopied that of both secretory RNases (Fig. 3f–g and Extended Data Fig. 2h–j). These results show that serum RNase A family members can mask the effects of RNASE4 deficiency, and that endogenous and exogenous RNase A family members contribute to lysosomal RNA degradation in cultured cells.

### Specific RNAs are targeted for lysosomal degradation by autophagy

Cells deficient for RNASET2, PLD3, and RNASE4 (henceforth referred to as triple-KO or tKO cells) and grown without serum accumulate ∼100-times more RNA in lysosomes than WT cells (Fig. 4a–b and Extended Data Fig. 3a). Interestingly, mTORC1 inhibition in these tKO cells cultured in the absence of serum did not substantially further increase lysosomal RNA amounts, which indicated that lysosomes in these cells cannot acquire much more RNA and that most of the RNAs in untreated and Torin1 treated tKO lysosomes were basally targeted for degradation.

**Fig. 4:**
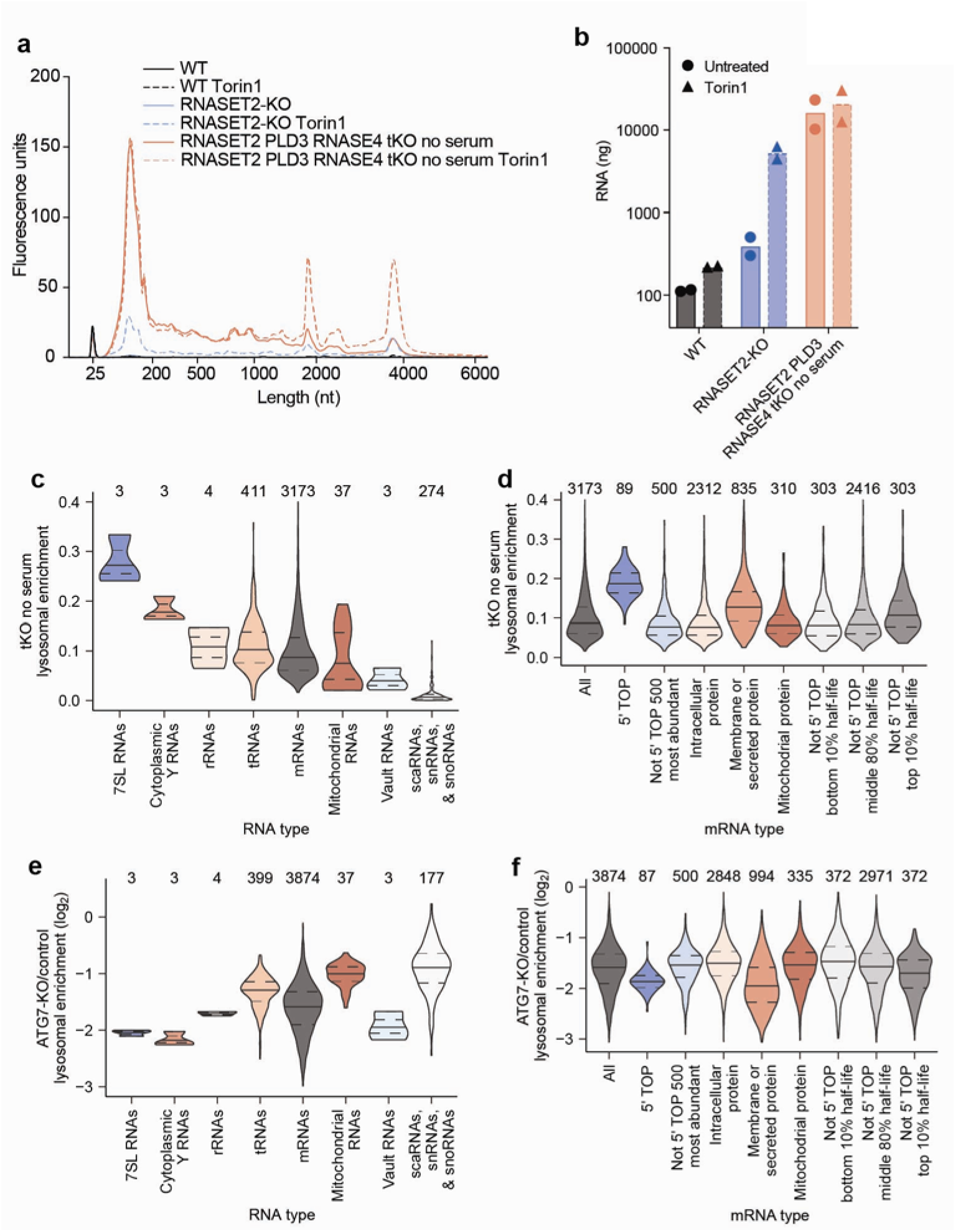
Specific RNAs are targeted for lysosomal degradation by autophagy. **a**, Effect of RNASET2, PLD3, RNASE4, serum and Torin1 on lysosomal RNA accumulation. Lysosomal RNAs from equal proportions of immunoprecipitates prepared for Extended Data Fig. 3A were purified and resolved on a bioanalyzer. The peak at 25 nt is a marker. Torin1 treatment was 250 nM for 5 h. **b**, Quantification of lysosomal RNAs. Lysosomal RNA amounts from the purified lysosomes of Extended Data Fig. 3A were quantified by bioanalyzer. **c**, Lysosomal enrichments for different RNA types in tKO cells grown without serum. The number above each violin plot indicates how many unique RNAs were considered in that plot. **d**, Lysosomal enrichments for different mRNA types in tKO cells grown without serum. Protein localization data is from the Human Protein Atlas^39^, mitochondrial proteins were defined by Mitocarta3^40^, and half-life data are from Agarwal and Kelly^41^. The number above each violin plot indicates how many unique mRNAs were considered in that plot. **e**, Effect of ATG7 KO on lysosomal enrichment for different RNA types. This panel is as in (**c**) except cells were ATG7 KO in addition to RNASET2, PLD3, and RNASE4 KO. **f**, Effect of ATG7 KO on lysosomal enrichment for different mRNA types. This panel is as in (**d**) except cells were ATG7 KO in addition to RNASET2, PLD3, and RNASE4 KO.

To identify lysosomally degraded RNAs in an unbiased manner, we performed quantitative RNA sequencing using Thermostable Group II Intron Reverse Transcriptase (TGIRT) on RNAs isolated from lysosomes from WT cells, RNASET2-KO cells, and tKO cells cultured in the absence of serum^38^. For all genotypes, we prepared cells treated without or with Torin1. To calculate lysosomal and whole-cell read counts for a given treatment, RNA sequencing reads were normalized using RNA spike-ins and lysosomal capture efficiencies estimated by LAMP2 amounts as assessed by immunoblotting. To calculate enrichment values, the lysosomal read counts were divided by whole-cell read counts.

Overall, similar types of RNAs were enriched within lysosomes across different genotypes and treatments, as demonstrated by moderate-or-better correlations between the enrichment values (Extended Data Fig. 3b; Spearman correlation coefficients (*R*_s_ values) ranging from 0.36–0.82). The population of RNAs enriched in untreated WT lysosomes correlated the least well with those from other conditions, perhaps because these lysosomes contained very little RNA, and thus a substantial fraction of the signal presumably came from mitochondrial contamination of purified lysosomes (Extended Data Fig. 3f). Lysosomal enrichments from tKO and tKO Torin1-treated cells had the highest correlation, which is consistent with the minimal effect of Torin1 treatment on bulk RNA accumulation in lysosomes in these cells. In contrast, lysosomal enrichments from RNASET2-KO cells treated with Torin1 correlated less well with those from tKO cells, suggesting that the RNAs targeted for degradation under Torin1 treatment are a similar but distinct set of RNAs.

Examination of the RNAs enriched within lysosomes showed that specific types of RNAs behaved differently. 7SL Signal Recognition Particle (SRP) RNAs were the most enriched type of RNA across most genotypes and treatments, followed by cytoplasmic Y RNAs. Perhaps as expected, nuclear RNAs such as scaRNA, snRNAs, and snoRNAs were relatively depleted from lysosomes (Fig. 4c, Extended Data Fig. 3c–g, and Supplementary Table 3).

In general, tRNAs were identified within lysosomes, but there was variability in the species of tRNAs enriched across genotypes and treatments (Extended Data Fig. 3h–n). tRNAs for negatively charged amino acids (glutamic acid and aspartic acid), were often more highly enriched (Extended Data Fig. 3i–n). In addition, in tKO cells cultured in the absence of serum, lysosomal tRNA enrichment correlated modestly with the hydropathicity^42^ of the amino acid carried by the tRNA (Extended Data Fig. 3h; *R*_s_ values ranging from 0.37–0.39). What accounts for the specificity of tRNAs by their cognate amino acid identity is an interesting area for future research.

We also examined whether specific types of mRNAs were targeted for lysosomal degradation. Many of the mRNAs that were enriched in lysosomes encode ribosomal proteins and translation factors, a trait of 5′ terminal oligopyrimidine (5′ TOP) mRNAs, which are characterized by the presence of 5–15 pyrimidines at their 5′ end^43^. La-related protein 1 (LARP1) binds to the 5′ TOP sequence and represses the translation of these transcripts in nutrient-deprived states^44^. Classically defined 5′ TOP mRNAs^45^ were lysosomally enriched across all genotypes and treatments (Fig. 4d and Extended Data Fig. 4a–e). Although these transcripts are very abundant, their abundance did not appear to cause their lysosomal enrichment because the 500 most abundant non-5′- TOP transcripts behaved like bulk mRNAs and not 5′ TOP mRNAs (Fig. 4d and Extended Data Fig. 4a–e). When mRNAs were binned by TOPscore, which reports on the strength of the 5′ TOP motif for each mRNA^45^, mRNAs with higher TOPscores were more lysosomally enriched across all genotypes and treatments (Extended Data Fig. 4f–k). Thus, the strength of the 5′ TOP motif of an mRNA serves as a good predictor of the extent to which it undergoes lysosomal degradation.

In addition to 5′ TOP mRNAs, mRNAs encoding membrane and secretory proteins (defined by the human Protein Atlas^39^) were enriched in lysosomes (Fig. 4d and Extended Data Fig. 4a–e). This was not the case for mRNAs of mitochondrial proteins (defined by Mitocarta3^40^), which implied that these mRNAs enter lysosomes because the autophagosome forms at the endoplasmic reticulum (ER) or from the selective degradation of the ER by ER-phagy^46,47^.

Given that 5′ TOP mRNAs are some of the most stable transcripts in the cell, we were curious if other long-lived transcripts were also lysosomally degraded. We compared a cell-line-independent mRNA half-life metric^41^ with the lysosomal enrichment score and found that transcripts with long half-lives were more likely to accumulate in lysosomes (Fig. 4d and Extended Data Fig. 4a–e). Interestingly, across all genotypes, this effect was more pronounced upon mTORC1 inhibition of cells (Fig. 4d and Extended Data Fig. 4a–e).

To test whether the lysosomal entry of these RNAs requires autophagy, we created tKO cells also deficient for the autophagy factors ATG7 or FIP200 and cultured them in the absence of serum (Extended Data Fig. 5a). Although all types of RNAs had some dependence on autophagy for lysosomal entry, some transcripts were more dependent on autophagy than others, implying that selective mechanisms mediate their lysosomal entry. In particular, many lysosomally enriched RNAs like 7SL RNAs, Y RNAs, 5′ TOP mRNAs, membrane-protein mRNAs, secretory-protein mRNAs, and long-lived mRNAs were more sensitive to the loss of autophagy than average mRNAs, rRNAs, mitochondrial RNAs, and small nuclear RNAs (Fig. 4c-f, Extended Data Fig. 5b–e, and Supplementary Table 4). Interestingly, vault RNAs were also highly sensitive to loss of the autophagy machinery despite not being particularly enriched within lysosomes (Fig. 4c, 4e, Extended Data Fig. 5b, and Supplementary Table 4). This is consistent with work identifying the vault as a substrate for autophagy and implies low flux but high specificity of the targeting of vault RNAs for degradation in lysosomes^48^. Together, our sequencing results identify specific RNAs that are preferentially degraded within the lysosome and establish that the autophagy machinery mediates both the lysosomal entry of all RNA types and confers specificity to the RNAs degraded by lysosomes.

### Lysosomally degraded RNAs have autophagy-dependent changes in stability

To determine the impact of lysosomal RNA degradation on the overall stability of RNAs within cells, we measured RNA half-lives in WT and ATG7-KO cells with and without autophagy activation caused by mTORC1 inhibition. We could not use approach-to-steady-state measurements for RNA stability, because these measurements rely on RNA synthesis rates being constant which is not the case in cells treated with an mTORC1 inhibitor^49^. Therefore, we set out to perform pulse-chase measurements using the uridine analog 5-ethynyl-uridine (5EU).

We initially collected RNA samples as soon as 0.5 h after labeling. However, we found that incorporation of 5EU continued to be detected up to 2 h after a short pulse of 5EU, consistent with previous findings^50^. This continued incorporation was observed in RNAs of varied stabilities. For example, normalized 5EU-containing reads mapping to the short-lived *MYC* mRNA continued to increase for 1 h post labeling, while normalized 5EU-containing reads mapping to the long-lived 18S rRNA increased for approximately 2 h post labeling (Extended Data Fig. 6a–b). This continued incorporation was presumably a consequence of 5EU retention in the cell after being converted into a nucleoside triphosphate, which reduced the efficacy of the chase.

To avoid acquiring data during the period of continued 5EU incorporation, we modified the pulse-chase protocol to pulse-label the cells for 20 min and then chase for 2 h before beginning the time course. We then collected RNA at 0, 1, 2, 4, 8, and 28 h, spiked-in a known amount 5EU-labeled RNA standards into each sample, biotinylated the labeled RNA, fragmented the RNA (a step designed to reduce the effect of incorporating any lingering 5EU), isolated the biotinylated RNA fragments by streptavidin pulldown, reverse transcribed biotinylated RNA using TGIRT, and amplified and sequenced the complementary DNA. Normalized reads were fit for decay rates to provide estimates of RNA half-lives (Fig. 5a). Our mRNA half-lives from WT whole-cell samples correlated well with previously reported mRNA half-lives^41^ (Extended Data Fig. 6c and table S5, *R*_s_ = 0.87), which confirmed the accuracy of our approach.

**Fig. 5:**
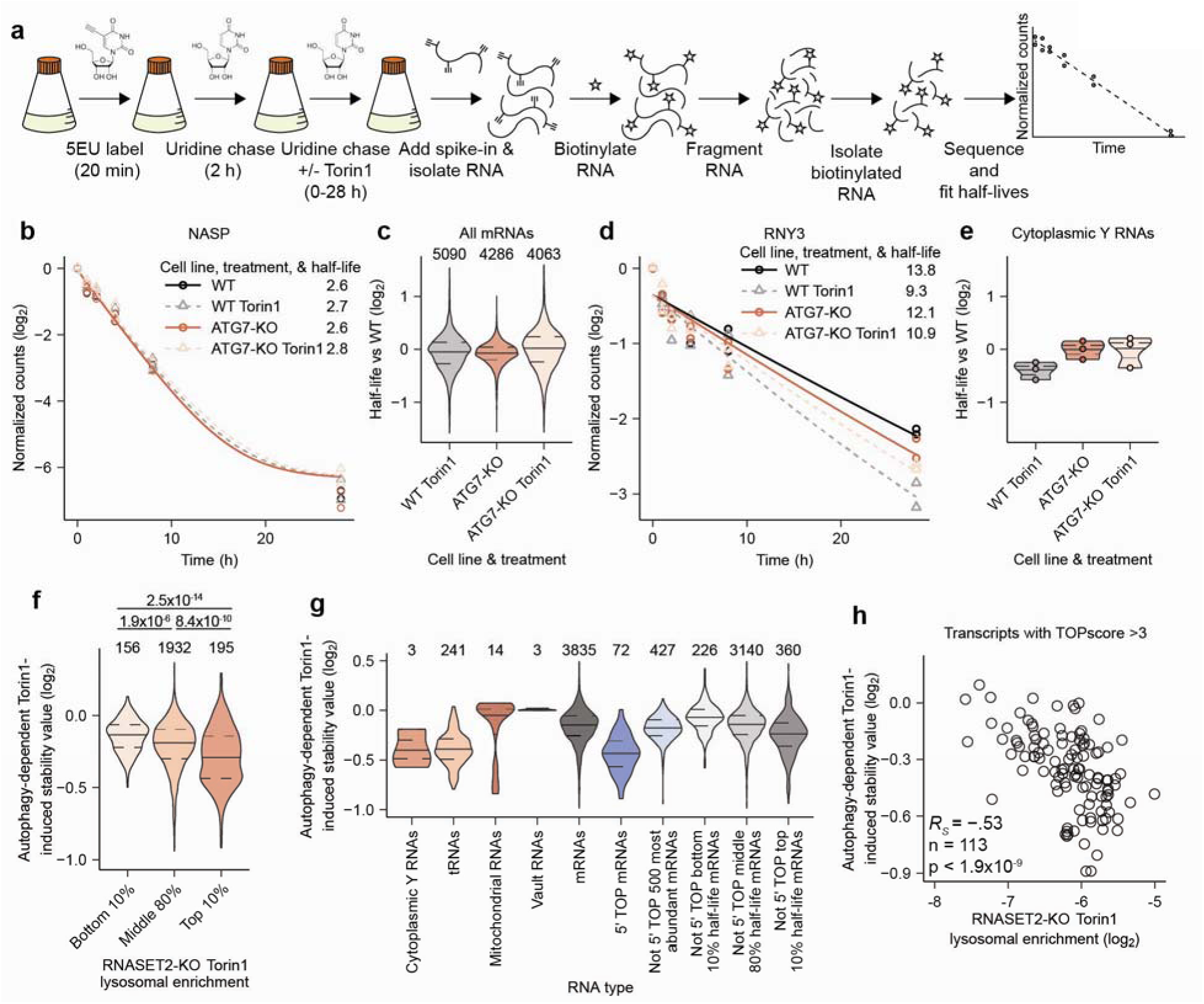
Lysosomally degraded RNAs have autophagy-dependent changes in stability. **a**, Schematic of 5EU labeling strategy for RNA half-life determination. **b**, Representative plot of effects of autophagy and Torin1 on mRNA stability. For each of the indicated genotypes and treatments, the lines represent decay equations determined by the average of decay rates and the average of an initial constant value established from biological replicates. **c**, Effects of autophagy and Torin1 on mRNA stabilities. The number above each violin plot indicates how many unique mRNAs were considered in that plot. **d**, Representative plot of effects of autophagy and Torin1 on Y RNA stability. Otherwise, this panel is as in (**b**). **e**, Effects of autophagy and Torin1 on Y RNA stabilities. Otherwise, this panel is as in (**c**). **f**, Autophagy-dependent Torin1-induced stability values grouped by lysosomal enrichment in RNASET2-KO cells treated with Torin1. The autophagy-dependent Torin1-induced stability value is defined by Torin1-induced changes in RNA half-lives in WT cells (WT Torin1 half-life/ WT half-life) divided by Torin1-induced changes in RNA half-lives in ATG7-KO cells (ATG7-KO Torin1 half-life/ ATG7-KO half-life). The number above each violin plot indicates how many unique RNAs were considered in that plot. The number between violin plots indicate the *P* value calculated using a Wilcoxon rank sum test. **g**, Autophagy-dependent Torin1-induced stability values grouped by RNA type. Otherwise, this panel is as in (**f**). **h**, Relationship between autophagy-dependent Torin1-induced stability values and lysosomal enrichments in RNASET2-KO cells treated with Torin1 for transcripts with TOPscores^45^ greater than 3.

With this method in hand, we determined RNA stabilities in WT and ATG7-KO cells with and without mTORC1 inhibition. Global RNA half-lives from these four conditions correlated well, indicating that large global changes in RNA stability did not occur following loss of autophagy (Extended Data Fig. 6d). For some types of RNAs, we were unable to calculate half-lives. For example, some did not fit single-exponential decay kinetics, which precluded accurate half-life calculations, and thus were excluded from our analyses (7SL RNAs, rRNAs, and many snRNAs, snoRNAs, and scaRNAs). Nonetheless, we were able to calculate half-lives for many mRNAs, tRNAs, Y RNAs, vault RNAs and mitochondrial RNAs.

To identify the role of autophagy in regulating RNA stability, we compared how Torin1 changed RNA half-lives in WT cells (WT Torin1 half-life/WT half-life) versus ATG7-KO cells (ATG7-KO Torin1 half-life/ATG7-KO half-life) by dividing the former ratio by the latter. By first normalizing within the same genotype, we accounted for any differences in base-line stability that might have resulted from variations in clonal cell lines. Then, by comparing between genotypes, we isolated changes in stability that were a result of autophagy from any broader effects of mTORC1 inhibition. We defined this metric as the autophagy-dependent Torin1-induced stability value. An autophagy-dependent Torin1-induced stability value less than 1.0 indicates that autophagy decreases the half-life of an RNA following mTORC1 inhibition. An example mRNA with a value near 1.0 is in Fig. 5b. This is representative of bulk mRNAs, for which half-lives were largely unperturbed across all four conditions (Fig. 5c). An example Y RNA with a value lower than 1.0 is in Fig. 5d. This is representative of all cytoplasmic Y RNAs, for which half-lives decreased following Torin1 treatment of WT cells, whereas Torin1 had little effect on the half-lives in ATG7-KO cells (Fig. 5e).

The autophagy-dependent Torin1-induced stability values of RNAs mostly corresponded weakly with their lysosomal enrichments (Extended Data Fig. 6e). This weak relationship was perhaps expected because only RNAs with high lysosomal enrichment should have a strong autophagy dependence for their stability. Of note, the strongest correlation between the lysosomal enrichment of RNAs and their autophagy-dependent Torin1-induced stability changes was in RNASET2-KO cells treated with Torin1 (Extended Data Fig. 6e–f). We suspect that this was because the enrichments for RNASET2-KO cells treated with Torin1 best reflected which RNAs are normally targeted for lysosomal RNA degradation following Torin1 treatment, whereas the tKO enrichments were more reflective of RNAs targeted for lysosomal degradation under basal conditions.

When RNAs were binned by lysosomal enrichment in Torin1-treated RNASET2-KO cells, the autophagy-dependent Torin1-induced stability values were lower for those that were more lysosomal (Fig. 5f). This relationship indicated that the lysosomal enrichments we identified are predictive of the impact of autophagy on RNA stability.

Furthermore, when examined by RNA type, many of those that we previously identified as lysosomally enriched, such as cytoplasmic Y RNAs, tRNAs, 5′ TOP mRNAs, and long-lived mRNAs, had correspondingly lower autophagy-dependent Torin1-induced stability values (Fig. 5g, and Extended Data Fig. 6g–h). mRNAs encoding membrane or secretory proteins that we identified as lysosomal however, did not have decreased autophagy-dependent Torin1-induced stability values (Extended Data Fig. 6i). This may have been because we were unable to calculate half-lives for many of the lysosomal membrane or secretory protein mRNAs (Extended Data Fig. 6j), which was explained by their lower 5EU-labeled read counts (Extended Data Fig. 6k–l).

5′ TOP mRNAs had low autophagy-dependent Torin1-induced stability values in a different manner from other RNAs. For these RNAs Torin1 treatment did not destabilize RNAs in WT cells, but Torin1 treatment caused their stabilization in cells lacking autophagy. Therefore, autophagy prevents the excess stabilization of 5′ TOP RNAs following mTORC1 inhibition (Extended Data Fig. 6m–n). As for lysosomal enrichment, we found that TOPscore also predicted autophagy-dependent Torin1-induced stability values (Extended Data Fig. 6o, and Supplementary Table 5). Indeed, lysosomal enrichment of mRNAs with a TOPscore greater than 3.0 moderately anticorrelated with autophagy-dependent Torin1-induced stability values (Fig. 5h, *R*_s_ = -0.53). Together, our measurements uncovered a role for lysosomal RNA degradation in regulating RNA stability.

### The Alu Domain of SRP RNA is sufficient for lysosomal entry in a manner that is dependent upon SRP9 and SRP14 binding

The 7SL SRP RNA is one of the most lysosomally enriched RNAs, and its lysosomal entry is highly autophagy-dependent (Fig. 4c, 4e, and Extended Data Fig. 5b). The 7SL RNA is the backbone of the SRP upon which the six SRP proteins assemble, and this complex is necessary for the translocation of nascent polypeptides into the ER^51^. The SRP RNA is divided into two domains, Alu and S. The Alu domain binds a heterodimer of SRP9 and SRP14 (SRP9/14) and enables the elongation-arrest activity of the SRP, while the S domain binds the four remaining SRP proteins and targets the SRP-bound translating ribosome to the ER^51^.

Analysis of truncation mutants of the SRP RNA in tKO cells cultured without serum, indicated that the Alu domain was sufficient to recapitulate the lysosomal entry of the full-length RNA (Fig. 6a–b). Indeed, the long Alu domain was more lysosomally enriched than full-length SRP RNA. Therefore, our subsequent experiments used this domain.

**Fig. 6:**
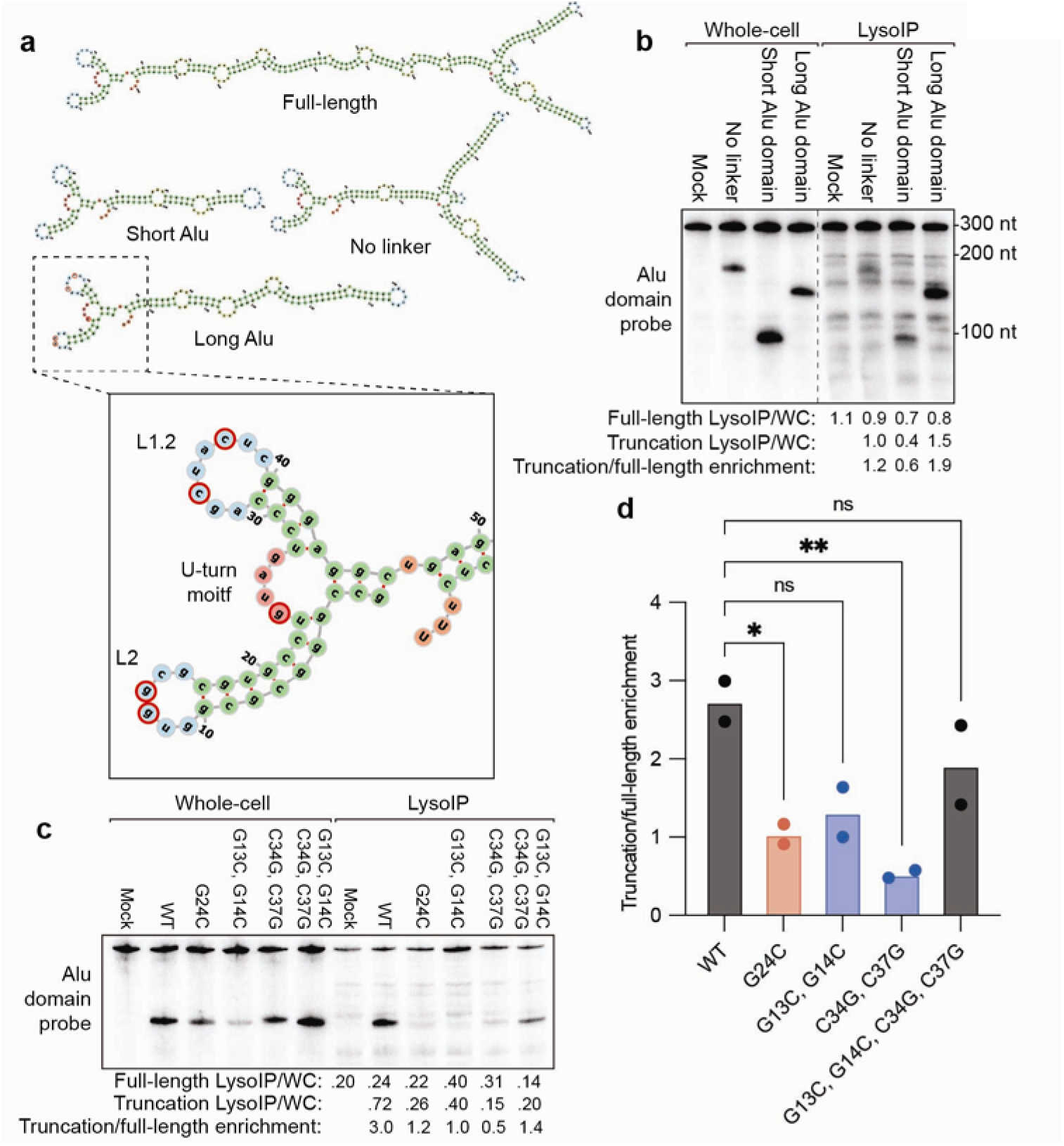
The Alu Domain of SRP RNA is sufficient for lysosomal entry in a manner that is dependent upon SRP9 and SRP14 binding. **a**, Schematic of 7SL SRP RNA and truncations. Images were made using forna based on the secondary structure reported in Zwieb *et al.*^54^. Mutated residues for (**c**) and (**d**) are highlighted by red circles in the inset. **b**, Sufficiency of the long Alu domain for lysosomal localization. RNASET2, PLD3, RNASE4 tKO HEK293T cells stably expressing TMEM192-3x-HA grown in serum-free media were transfected with plasmids encoding the indicated SRP RNA truncations. Five days after transfection, equal cell numbers were subjected to the LysoIP protocol, and whole-cell and lysosomal RNA were analyzed by northern blot. Shown is a representative blot. **c**, Effect of mutants that disrupt SPR9/14 binding to the SRP Alu Domain on lysosomal localization. The experiment was performed as in (**b**), except that LysoIPs and RNA collection were performed four days after transfection. Shown is a representative RNA blot. **d**, Quantification of (**c**) and a replicate experiment. Significance was evaluated by one-way ANOVA (ns, *P* value >.05; *, *P* value <.05; **, *P* value <.01).

Because the Alu domain of the SRP RNA binds SRP9/14, we tested if SRP9/14 binding is necessary for the lysosomal entry of the SRP RNA by introducing point mutations known to disrupt SRP9/14 binding. The first mutant, G24C in the U-turn motif of the SRP RNA, reduces SRP9/14 binding by 50-fold^52^. In addition, mutations in the L2 and L1.2 loops prevent the ability of the loops to pair with each other, which also disrupts SRP9/14 binding^53^. Moreover, combining the mutations in L2 and L1.2 restores base pairing and SRP9/14 binding^53^. All mutations predicted to reduce the ability of the Alu domain to bind SRP9/14 also reduced lysosomal entry of the truncated SRP RNA (Fig. 6c–d). Thus, binding of SRP9 and SRP14 to the SRP RNA is necessary for its efficient lysosomal entry.

### 5’ TOP motifs target transcripts for LARP1-dependent lysosomal degradation

Having found that 5′ TOP mRNAs were lysosomally degraded in an autophagy-dependent manner (Fig. 4d, 4f, and Extended Data Fig. 4 and 5c–e), we wanted to examine whether the 5′ TOP motif is required to specify the lysosomal entry of this mRNA type. To do so, we constructed reporters that varied at their 5′-terminal nucleotides and examined their localization in tKO cells cultured without serum. The reporters were based on the 5′ UTR of the elongation factor 1 alpha (EF1alpha) mRNA, which was placed upstream of a green fluorescent protein (GFP) coding sequence (Fig. 7a). This 5′ UTR contains a 5′ TOP motif at its first seven nucleotides. Reporter variants containing either this motif or another strong 5′ TOP motif were ∼3-fold more lysosomal than variants that did not contain a 5′ TOP motif (Fig. 7b). Thus, the 5′ TOP motif is required to promote the lysosomal entry of mRNAs.

**Fig. 7:**
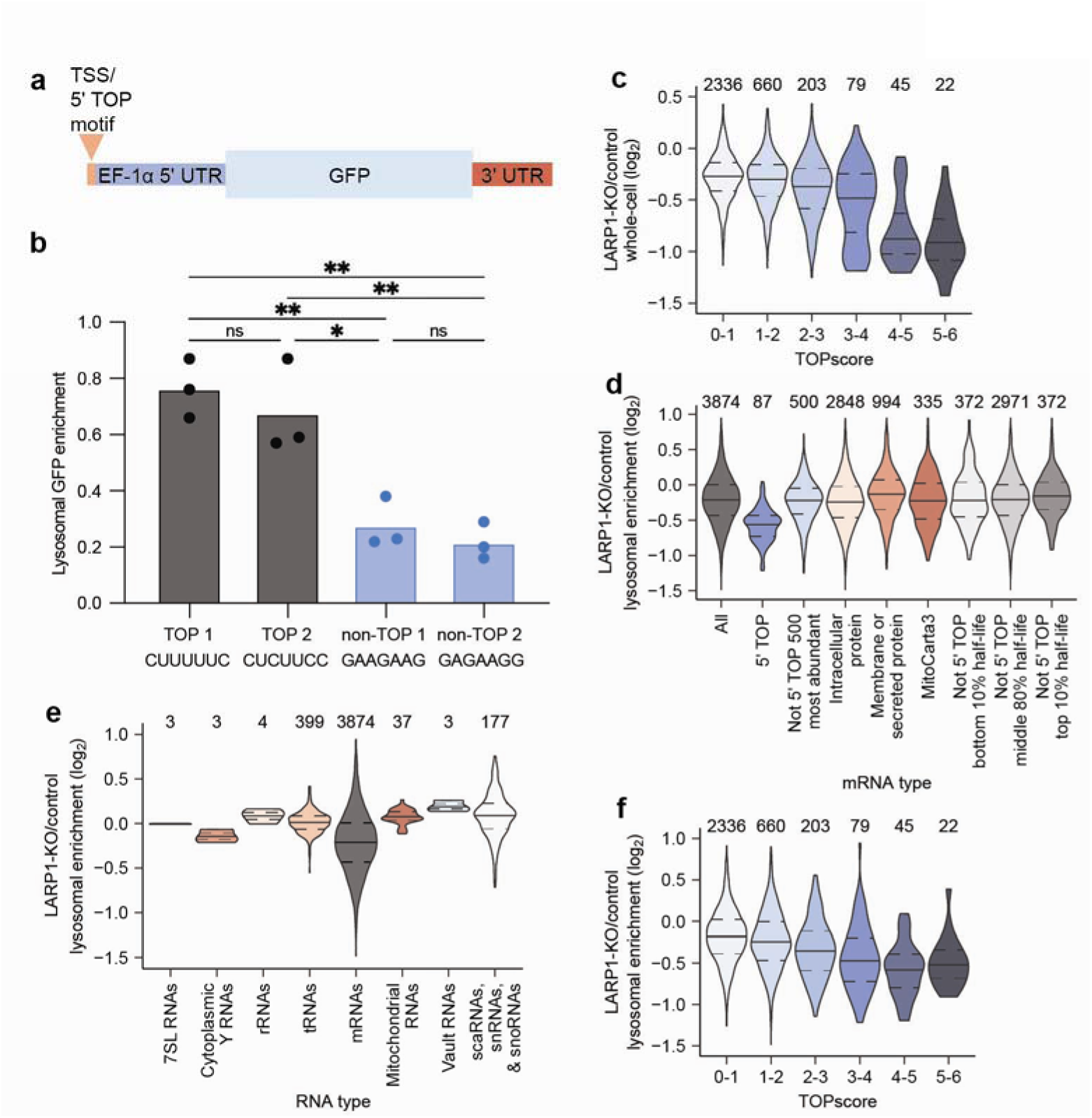
5′ TOP motifs target transcripts for LARP1-dependent lysosomal degradation. **a**, Schematic of the GFP reporter. Variants differ only in their first seven nucleotides. **b**, Effect of 5′ TOP motif on lysosomal targeting of the GFP reporter. RNASET2, PLD3, RNASE4 tKO HEK293T cells stably expressing TMEM192-3x-HA grown in serum-free media were transfected with plasmids encoding the indicated GFP reporter transcripts. Four days after transfection, equal cell numbers were subjected to the LysoIP protocol, and whole-cell and lysosomal RNA were analyzed by quantitative PCR. The amount of GFP mRNA was normalized to that of the 18S rRNA in both whole-cell and lysosomal fractions. 3 independent transfections were done for each reporter. Significance was evaluated by one-way ANOVA (ns, *P* value >.05; *, *P* value <.05; **, *P* value <.01). **c**, Effect of LARP1 KO on whole-cell 5′ TOP stability. The number above each violin plot indicates how many unique RNAs were considered in that plot. **d**, Effect of LARP1 KO on lysosomal enrichment for different mRNA types. Protein localization data is from the Human Protein Atlas^39^, mitochondrial proteins were defined by Mitocarta3^40^, and half-life data are from Agarwal and Kelly^41^. The number above each violin plot indicates how many unique mRNAs were considered in that plot. **e**, Effect of LARP1 KO on lysosomal enrichment for different RNA types. The number above each violin plot indicates how many unique RNAs were considered in that plot. **f**, Effect of LARP1 KO on lysosomal enrichment as a function of TOPscore^45^. The number above each violin plot indicates how many unique RNAs were considered in that plot.

Because 5′ TOP motifs are bound by LARP1, we wondered whether LARP1 influences the lysosomal entry of 5′ TOP mRNAs. To investigate this possibility, we knocked out LARP1 in the tKO cells and performed quantitative RNA sequencing on lysosomal and whole-cell fractions of the cells cultured in the absence of serum (fig. S5A). As reported in other contexts, loss of LARP1 led to a decrease in the steady-state levels of 5′ TOP mRNAs (Fig. 7c and Supplementary Table 4)^55^. With respect to localization, loss of LARP1 substantially decreased lysosomal enrichment of mRNAs with 5′ TOP motifs, with negligible impact on lysosomal entry of other RNAs (Fig. 7d–e and Supplementary Table 4). Furthermore, the dependency of the mRNA on LARP1 for lysosomal entry scaled with the strength of the 5′ TOP motif, in that mRNAs with higher TOPscores were more dependent on LARP1 for lysosomal enrichment (Fig. 7f and Supplementary Table 4). Therefore, LARP1 helps facilitate the trafficking of 5′ TOP mRNAs to lysosomes for degradation.

## Discussion

We identified RNASET2, PLD3, and both endogenous and exogenous RNase A family members as lysosomal RNases. These RNases collectively degrade endogenous RNAs that are specifically targeted to the lysosome by autophagy. In particular, the SRP RNAs, Y RNAs, 5′ TOP mRNAs, long-lived mRNAs, membrane mRNAs, and secretory mRNAs are preferentially targeted to lysosomes for degradation. Metabolic labeling of RNAs to calculate RNA half-lives demonstrated that RNAs targeted for degradation within the lysosome tend to be destabilized by autophagy. We defined two sequence elements that are sufficient to increase lysosomal targeting: the Alu domain of the SRP RNA in a manner that depends upon SRP9/14 binding, and 5′ TOP motifs in a manner that depends upon LARP1. In sum, we establish lysosomal RNA degradation via autophagy as an additional mechanism of post-transcriptional gene regulation.

Why have cells evolved mechanisms to degrade RNAs lysosomally as opposed to using canonical RNA decay mechanisms? One reason could be that the lysosome is better suited for the degradation of highly structured RNAs or ribonucleoprotein (RNP) complexes, such as the SRP RNA and Y RNAs we identified as particularly lysosomally enriched. The lysosome is also host to many proteases that could act in tandem with RNases to degrade RNPs ^3^. RNASET2 preferentially cleaves between a purine and uridine, RNase A cleaves after pyrimidines, and PLD3 is a 5′ to 3′ exonuclease that can degrade the cleavage products created by RNASET2 or RNase A^31,56,57^. We also hypothesize there remains at least one unidentified lysosomal RNase, because we observed residual degradation patterns in the lysosomes of tKO cells grown without serum in SRP RNA northerns (Fig. 6b–c). Perhaps this combination of lysosomal RNases (with potentially others in different cell types) and proteases collaborate to detangle and destroy complex RNA or RNP structures more efficiently than the mechanisms that operate in cytoplasm.

Another possibility is that degrading heavily modified RNAs, such as tRNAs and rRNAs, within the lysosome, which uses a unique degradative chemistry, might be beneficial.

Cytoplasmic decay produces nucleotides with a 5′ phosphate, whereas lysosomal decay produces nucleotides with a 3′ phosphate ^19^. Nucleotides with 3′ phosphates must be fully dephosphorylated to nucleosides before they can have a 5′ phosphate added and subsequently reincorporated into RNAs, whereas nucleotides with a 5′ phosphate can simply have two phosphates added before reincorporation into new RNAs^58^. Thus, the 3′ phosphate removal and 5′ phosphate addition, which lysosomally derived nucleosides must undergo, might provide additional opportunities to discriminate between modified and unmodified nucleosides/nucleotides and thereby reduce aberrant incorporation of modified nucleotides into nascent RNAs.

Lastly, lysosomal RNA degradation could serve as a way of degrading RNAs that are not degradable by canonical decay mechanisms. One such example may be 5′ TOP mRNAs. Upon mTORC1 inhibition, LARP1 binds the 5′ cap of 5′ TOP mRNAs to prevent their translation (and possibly decapping) while also binding PABP (poly (A)- binding protein) and the 3′ end of their poly(A)-tails, which could block deadenylation^44,45,59–62^. Thus, these LARP1-bound mRNAs might be resistant to deadenylation, decapping, or any RNA decay mechanism that requires translation, such as no-go decay, non-stop decay, or nonsense-mediated decay. Perhaps autophagic degradation of 5′ TOP mRNAs circumvents these limitations so that 5′ TOP mRNAs can be degraded without requiring deadenylation and decapping. This model is also consistent with the dual roles for LARP1 that we observed: stabilization of 5′ TOP mRNAs under basal conditions while also specifying the preferential degradation of 5′ TOP mRNAs by autophagy.

The variability in the rates of basal and stress-activated autophagy across mammalian cell types has become increasingly appreciated, as has the observation that dysfunctional autophagy can contribute to many pathologies, including neurodegeneration^63,64^. The methods and insights developed here might facilitate the study of lysosomal RNA degradation *in vivo*, so as to define the types and extents of RNAs degraded via the lysosome in different cell types and disease states.

## Methods

### Cell culture

HEK293T cells for virus production were cultured in DMEM + GlutaMAX (Gibco 10569- 010) supplemented with 10% FBS (Gibco 10438-026) and 1% Pen-strep (Gibco 15140- 122) at 37°C with 5% CO_2_. HEK293T cells for all other work were cultured in Freestyle 293 (Gibco 12338-018) supplemented with 1% or 0% FBS (Gibco 10438-026) and .5% Pen-strep (Gibco 15140122). To adapt cells to freestyle media, cells were passaged from DMEM media, into progressive mixes of DMEM to Freestyle (50:50, 25:75, 10:90, 0:100). At the point that cells were in 100% Freestyle, they were grown in shaker flasks and grown in an infors shaker at 37°C shaking at 125 RPM, with 8% CO_2_, and 80% humidity.

### KO cell lines

Clonal KO lines derived from HEK293T cells were generated by transiently transfecting cells with plasmids encoding Cas9, a gRNA, and either a fluorescent protein, or selectable marker. The backbones used were pX458 (Addgene #48138), pX459 (Addgene #48139), or pEF60 and single-cell sorting cells GFP positive cells, puromycin resistant cells, or mScarlett positive cells respectively. Most clonal lines were genotyped by PCR and sequencing for mutations that would lead to frame-shift mutations. In the instance of ATG7, two guides were used and clones with mutations that would result in removal cysteine 572, which is necessary for canonical ATG7 activity, were selected by PCR and sequencing. In the instance of RNASE4/ANG, two guides were used and PCR was used to find clones with homozygous deletion of the regions between the two guides. Often genotyping primers included T3 and T7 sequences such that genotyping PCRs could be barcoded in a second PCR reaction to enable multiplex genotyping by NGS. Primers used for cloning and genotyping are below.

RNASET2 KO clonal lines were transfected with pGJR002 (RNASET2 sgRNA 1 in pX459) cloned with oGJR001:CACCGTCCAGGCCCGATAAAAGTGA and oGJR004:AAACTCACTTTTATCGGGCCTGGAC or pGJR055(RNASET2 sgRNA 4 in pX458) cloned with oGJR154:CACCGCCGGATTACTGGACAATACA and oGJR156:AAACTGTATTGTCCAGTAATCCGGC.

RNASET2 sgRNA 1 clones were genotyped with oGJR098:GAACTCTGACTCCTCTTACCTGC and oGJR099:GATCTGCACAGGTCTGGCATG.

RNASET2 sgRNA 4 clones were genotyped with oGJR166:GTATCTCGTTGGCATATGTTGC and oGJR167:GACCACATTCAAGCTAGGGTTC

PLD3 KO clonal lines were transfected with pGJR267(PLD3 sgRNA 2 in pEF60) cloned with oGJR696:CACCGACTTCCCCAATGCCTCCACG and oGJR698:AAACCGTGGAGGCATTGGGGAAGTC.

PLD3 g2 KO clones were genotyped with oGJR721:TAATACGACTCACTATAGGGGTATGGCTGATAGCATCCCC and oGJR722:CGATTAACCCTCACTAAAGGATTGTTGGTGAGGGTCCAGTAG

RNASE4 KO clonal lines were transfected with pGJR099 (RNASE4 sgRNA 2 in pX458) cloned with oGJR312:CACCGATCGCTGGTACATGCCATCC and oGJR314:AAACGGATGGCATGTACCAGCGATC.

RNASE4 g2 KO clones were genotyped with oGJR317:TAATACGACTCACTATAGGGATTCATTGCTTCTGCTTTTGCT and oGJR318:CGATTAACCCTCACTAAAGGCTTTGCATCATCAAGTTGCAGT

RNASE4 ANG dKO clonal lines were transfected with pGJR097 (Px458 with ANG g2) cloned with oGJR304:CACCGGCACTATGATGCCAAACCAC and oGJR306:AAACGTGGTTTGGCATCATAGTGCC or pGJR098 (RNASE4 sgRNA 1 in pX458) cloned with oGJR311:CACCGGCCGTTCTTGCATTGGATAT and oGJR313:AAACATATCCAATGCAAGAACGGCC

RNASE4 ANG dKO clonal lines were genotyped by 3 PCRs. Seeking clones with no signal for a PCR around ANG g2 using oGJR309:TAATACGACTCACTATAGGGGTTTTGTTGTTGGTCTTCGTGC and oGJR310:CGATTAACCCTCACTAAAGGAATGTGTTGATGTCTTTGCAGG, no signal for a PCR around RNASE4 g1 using oGJR315:TAATACGACTCACTATAGGGATGCAAAGACGGAAGATGACTT and oGJR316:CGATTAACCCTCACTAAAGGCCTGCAATCTGTGACCTTCACT, and a positive short amplicon signal for a PCR spanning ANG and RNASE4 using oGJR309:TAATACGACTCACTATAGGGGTTTTGTTGTTGGTCTTCGTGC and oGJR316:CGATTAACCCTCACTAAAGGCCTGCAATCTGTGACCTTCACT.

ATG7 KO clonal lines were transfected with both pGJR062 (PX458 with ATG7 sgRNA 1) and pGJR063 (PX458 with ATG7 sgRNA 2). pGJR062 was cloned with oGJR204:CACCGCTGGTCCAAGGTCCGGTCTC and oGJR208: AAACGAGACCGGACCTTGGACCAGC. pGJR063 was cloned with oGJR205: CACCGTACCATCAATTCCACGGCCA and oGJR209:

AAACTGGCCGTGGAATTGATGGTAC. ATG7 KO clones were genotyped for mutations that would result in at least removal cysteine 572 with oGJR221:TAATACGACTCACTATAGGGGAACGTTCTGCACACACCAA and oGJR213:CGATTAACCCTCACTAAAGGTAGCAAACTCACCCTTCTGGAT.

Polyclonal KO lines derived from HEK293T cells were generated by transduction with lentivirus produced from the lentiCRISPRv2 transfer plasmid (Addgene #163126) followed by selection with puromycin. Oligos used for cloning are below:

pGJR171 with AAVS1 g1 in lentiCRISPRv2 was cloned with oGJR1068:CACCGGGGGCCACTAGGGACAGGAT and oGJR1069:AAACATCCTGTCCCTAGTGGCCCCC

pGJR172 with NT g1 in lentiCRISPRv2 was cloned with oGJR1070:CACCGAATAGCTCAGAGGCCGAGG and oGJR1071:AAACCCTCGGCCTCTGAGCTATTC

pGJR162 with ATG7 g1 in lentiCRISPRv2 was cloned with oGJR204:CACCGCTGGTCCAAGGTCCGGTCTC and oGJR208:AAACGAGACCGGACCTTGGACCAGC

pGJR163 with ATG7 g2 in lentiCRISPRv2 was cloned with oGJR205:CACCGTACCATCAATTCCACGGCCA and oGJR209:AAACTGGCCGTGGAATTGATGGTAC

pGJR164 with FIP200 g1 in lentiCRISPRv2 was cloned with oGJR206:CACCGGATAGACAGTAGACGAATGC and oGJR210:AAACGCATTCGTCTACTGTCTATCC

pGJR165 with FIP200 g2 in lentiCRISPRv2 was cloned with oGJR207:CACCGAGAGGTTGAACTTGCGTTGA and oGJR211:AAACTCAACGCAAGTTCAACCTCTC

pGJR258 with PLD3 g1 in lentiCRISPRv2 was cloned with oGJR695:CACCGTAGCGGGTGTCATAGAACCG and oGJ697:AAACCGGTTCTATGACACCCGCTAC

pGJR259 with PLD3 g2 in lentiCRISPRv2 was cloned with oGJR696:CACCGACTTCCCCAATGCCTCCACG and oGJR698:AAACCGTGGAGGCATTGGGGAAGTC

pGJR337 with RNASE4 g1 in lentiCRISPRv2 was cloned with oGJR311:CACCGGCCGTTCTTGCATTGGATAT and oGJR313:AAACATATCCAATGCAAGAACGGCC

pGJR338 with RNASE4 g2 in lentiCRISPRv2 was cloned with oGJR312:CACCGATCGCTGGTACATGCCATCC and oGJR314:AAACGGATGGCATGTACCAGCGATC

pGJR386 with LARP1 g2 in lentiCRISPRv2 was cloned with oGJR1029:CACCGGGGTGCCGGCGCATGTAATG and oGJR1033:AAACCATTACATGCGCCGGCACCCC

pGJR387 with LARP1 g3 in lentiCRISPRv2 was cloned with oGJR1030:CACCGGATGAGGATTGCCAGCGAGG and oGJR1034:AAACCCTCGCTGGCAATCCTCATCC

#### CRISPR screen

A lentiviral library encoding 97,888 unique sgRNAs in a Cas9-containing vector targeting ∼20,000 genes and noncoding RNAs, including one AAVS1 safe harbor locus-targeting, 125 intergenic-targeting, and 250 nontargeting sgRNAs^29^ was prepared by transfecting HEK293T cells with a mixture containing 20 µg plasmid library, 3.62 µg pCMV-VSV-G, and 8.28 µg psPAX2, 76.8 µL Xtremegene-9 (Roche XTG9-RO) transfection reagent, and Opti-MEM (Gibco 31985062) media to a final volume of 1 mL. Cells were seeded at 15 x 10^6^ cells in a T175cm^2^ flask in 20 mL DMEM supplemented with 10% fetal bovine serum and 1% penicillin-streptomycin 30h pre-transfection, and media was changed to 20 mL virus production media (IMDM (Gibco 31980039) supplemented with 20% heat-inactivated fetal bovine serum) 7-8h pre-transfection.

Media was changed to 55 mL fresh virus production medium 16-17 h post-transfection, and the viral supernatant was harvested 48h post-transfection by 0.45 µm filtration and stored at -80°C. The virus was concentrated 20x with lenti-x (Takara bio 631231).

The screen infection was done as outlined in Adelmann *et al.*^65^ with slight modifications. 570 million HEK293T RNASET2-KO cells grown in Freestyle 293 with 1% serum were transduced with 2.5 mL 20x concentrated virus and 228 uL 10 mg/mL polybrene (MilliporeSigma™ TR1003G) in a final volume of 228 mL to achieve ∼2,000-fold library coverage after puromycin selection. Cells were plated in 5 million cells/well in 6-well dishes and spun at 37°C for 45 min at 1200 x g. The next day viral media was removed, cells were washed with 1x with PBS, and each well was replated into a 15 cm plate with Freestyle 293 with 1% serum. The next day the media was changed to include 1 ug/mL puromycin. Two Days later, 800 million cells were passaged again into 1 ug/mL puromycin containing media but this time into shaker flasks in suspension at 0.5 million cells/mL. Cells were continued to be cultured, always maintaining at least 500 million cells. On days 13 and 17 post infection, 250 and 300 million cells, respectively, that had been treated with 250 nM Torin1 the evening prior were stained with acridine orange and subject to FACS. Cells with top and bottom 10% of acridine orange signals (red-to-green ratio) were sorted into BSA coated tubes. Cells were pelleted and stored at - 20°C.

Genomic DNA (gDNA) was isolated from cell pellets using the Blood genomicPrep Mini Spin (Cytiva 28904264) or QIAmp DNA Blood Maxiprep (Qiagen 51104) kit and quantified using the Qubit dsDNA HS Assay Kit (Invitrogen Q32851). Test PCR amplifications containing 1, 3, or 6 µg gDNA in a 50 µL total reaction volume were performed for 28 cycles using ExTaq DNA Polymerase (Takara Bio RR001A) with the following primers:

Forward: 5′- AATGATACGGCGACCACCGAGATCTACACCCCACTGACGGGCACCGGA - 3′

Reverse: 5′- CAAGCAGAAGACGGCATACGAGATCnnnnnnTTTCTTGGGTAGTTTGCAGTTTT - 3′

Where “nnnnnn” indicates the demultiplexing barcode.

Reactions containing either 3 or 6 µg gDNA per 50 µL reaction were subsequently scaled up to sample ∼100 ug of gDNA.

Test and scaled PCR reactions were pooled, an aliquot was purified using the DNA Clean and Concentrator-5 kit (Zymo D4014), and samples were sequenced for 50 cycles on an Illumina HiSeq2500 using the following primers:

Read 1 sequencing primer: 5′- GTTGATAACGGACTAGCCTTATTTAAACTTGCTATGCTGTTTCCAGCATAGCTCTTA AAC - 3′

Index sequencing primer: 5′- TTTCAAGTTACGGTAAGCATATGATAGTCCATTTTAAAACATAATTTTAAAACTGCAA ACTACCCAAGAAA - 3′

Reads were trimmed and mapped using Bowtie v1.0.0^66^ and counted. gRNAs that targeted multiple genes were omitted from further analysis. gRNA enrichment and statistics were calculated with MAGeCK with the two timepoints as replicates^67^.

### Lentivirus production and infection

0.75 million HEK293T cells were plated in a well of a 6-well plate. The next day, cells were transfected with 0.15ug of vsvg plasmid (Addgene #8454), 0.9 ug of dvpr plasmid (Addgene #8455), and 1 ug of the transfer plasmid using x-tremegene 9 (Roche XTG9-RO). The following day, the media was changed. The day after that, media was collected and centrifuged at 500 × *g* for 10 min to pellet cell debris.

To infect cells, 1 million HEK293T cells were plated in a well of a 6-well plate with 2 uL 10 mg/mL polybrene (MilliporeSigma™ TR1003G), variable amounts of virus, and media to 2 mL. Then, plates were spun at 37°C for 45 min at 1200 x g and incubated at 37°C overnight in a tissue culture incubator. The next day, virus containing media was removed and replaced with fresh media. The following day, cells were passaged into media containing selection antibiotic (puromycin or blasticidin). To ensure selection went to completion, cells were passaged one more time in media containing selection antibiotic.

### LysoIP and MitoIP

Isolations of lysosomes and mitochondria were as before^22^ with some modifications. Cells were transduced with lentivirus to stably express either the 3xHA-Lyso tag (pLJC5-TMEM192-3xHA (Addgene #102930)), Control-Lyso tag (TMEM192-2xFlag (Addgene #102929)), or the 3xHA-Mito tag (pMXs-3xHA-EGFP-OMP25 (Addgene #83356)) ^22,24^. Cells expressing these constructs were washed in PBS and resuspended in 1 mL of KPBS. Whole-cell samples were taken from this resuspension. Cells were dounced 20 times in a 1mL dounce (VWR 89026-386 & 89026-398), spun for 2 min at 1,000 x g, and 750 uL of the supernatant was added to 100 uL prewashed anti-HA beads (Thermo Scientific 88837). Beads were then incubated at 4°C for 20 min with rocking. Beads were washed 4x with 1mL KPBS with changing of tubes after the 1^st^, 3^rd^, and 4^th^ washes. After the last wash the IP was split into 2 tubes, one for RNA extraction and one for protein. The RNA sample was resuspended in 240 uL KPBS + 125 mM Sucrose + 1 U/uL RNAse I (Invitrogen AM2295) (and potentially 1% Triton X-100 for some experiments) and incubated at 37°C with shaking at 400 rpm for 30 min. 750 uL Trizol LS (Invitrogen 10296010) was then mixed with the IP and the sample stored at -80°C until extraction. The protein fraction was resuspended in 50 uL lysis buffer and allowed to rock for 10 min at 4°C before spinning at 21,000 × g for 10 min at 4°C and retention of the supernatant.

### RNA isolation for lysosomal enrichment experiments

Samples were purified using Trizol LS (Invitrogen 10296010) and the RNA clean and concentrator (Zymo R1015 or R1017). Briefly, 750 uL Trizol LS was mixed with 250 uL of sample (if the sample was less than 250 uL, water was added to 250 uL). Samples were homogenized by pipetting and frozen at -80°C if the RNA purification was not done the same day. After thawing, 200 uL chloroform was added per 750 uL of Trizol LS. Samples were mixed by shaking for 15 s and incubated at room temperature for 3 min. Then, samples were centrifuged for 15 min at 12,000 x g at 4°C. 500 uL of the upper aqueous phase was transferred to a new tube, 500 uL of 100% ethanol was added to the sample, and they were mixed by pipetting. 700 uL of this sample was then transferred to zymo-spin IC or IICR column in a collection tube and centrifuged at 10,000 x g for 30 s. The flow through was discarded. Additional sample was transferred to the same column and centrifuged at 10,000 x g for 30 s. Flow through was again discarded. 400 uL of RNA prep buffer was added to the column and centrifuged at 10,000 x g for 30 s. Flow through was discarded. 700 uL of RNA wash buffer was added to the column and centrifuged at 10,000 x g for 30 s. Flow through was discarded. 400 uL of RNA wash buffer was added to the column and centrifuged at 10,000 x g for 30 s. Flow through was discarded. The column was transferred to a new tube for elution. 15 or 25 uL of water was added to the column matrix and the sample was centrifuged at 10,000 x g for 30 s.

### Cell lysis & western blotting

Cells and lysosomes were lysed with Triton X-100 lysis buffer (1% Triton X-100, 40 mM HEPES pH 7.4, 10 mM β-glycerol phosphate, 10 mM pyrophosphate, 2.5 mM MgCl2) and 1 tablet of EDTA-free protease inhibitor mini (Roche) per 10 mL buffer. Cells or lysosomes on beads were incubated with rotating at 4°C for 10+ min before centrifugation at 21,000 × g at 4°C for 10 min and retention of the supernatant. Samples were denatured by incubating in a buffer containing 2% SDS, 5% BME, 5% glycerol, and 50 mM Tris with .01% bromophenol blue and run on 4-20% Tris-Glycine gels (Invitrogen) with Tris glycine SDS running buffer (Invitrogen LC2675-4) buffer at 120V for 1.5-2 h and then transferred in a buffer of 10 mM CAPS pH 11 with 10% ethanol onto .45 um PVDF transfer membranes (Millipore IPVH0010). Membranes were then blocked in 5% milk in TBST for 1 h, and incubated in primary antibody diluted in 5% BSA in TBST with overnight rocking at 4°C. The following day, membranes were washed three times in TBST for 5 min, incubated in secondary antibodies diluted in 5% milk in TBST for 1 h, washed again three times in TBST for 5 min, incubated with Pierce ECL Western blotting substrate (Thermo scientific 32106) or SuperSignal West Atto Ultimate Sensitivity substrate (Thermo scientific A38554) and imaged with CL-X Posure Films (Thermo scientific 34091) and developed on an X-OMAT. The following antibodies were purchased from commercial sources and used at the indicated dilutions: LAMP2 (SCBT sc-18822) 1:1000, CSTC (SCBT sc-74590) 1:500, RNASET2 (SCBT sc-393729) 1:500, ANG (SCBT sc-74528) 1:100, CTSB (CST 31718S) 1:500, CTSD (CST2284S) 1:500, LC3B (CST 3868S) 1:500, GOLGA1 (CST 13192S) 1:1000, S6K (CST 9202S) 1:1000, CALR (CST 12238S) 1:1000, VDAC (CST 4661S) 1:1000, ATG7 (CST 2631S) 1:1000, FIP200 (CST 12436S) 1:1000, LARP1 (CST 70180S) 1:1000, Anti-mouse IgG HRP linked antibody (CST 7076S) 1:3000, Ant-rabbit IgG HRP linked antibody (CST 7074S) 1:3000, PLD3 (SIGMA HPA012800-100) 1:500, RNASE1 (Invitrogen PA5-78151) 1:500.

### Serum fractionation

500 mL Fetal bovine serum (10438-026 Gibco) was loaded onto a HiTrap SP XL column (Cytiva) equilibrated in a buffer containing 50 mM PBS pH 8.0 and eluted with a linear gradient of NaCl concentration to 2 M. Fractions were then desalted into PBS with an Amicon^®^ Ultra Centrifugal Filter, 3 kDa MWCO (Millipore sigma UFC9003) and filter sterilized before adding to cell culture media.

### Proteomics

In Low Protein Binding Microcentrifuge Tubes (Thermo Scientific 90410), samples were mixed 1:1 with 2X SDC buffer (2% SDC, 200 mM TEAB, 80 mM CAA, 20 mM TCEP), heated at 70°C for 25 min, and digested overnight with 1 ug trypsin/Lys-C mix at 37°C in a shaking incubator at 115 RPM. The following day, an additional 1 ug trypsin/Lys-C mix was added and digestion continued for 4 h at 37°C. Peptide digests were purified using stage tips as in Rappsilber *et al.* ^68^. Eluates were dried in a speed-vac concentrator and resuspended in 0.2% formic acid in MS-grade water.

LC-MS/MS data measurements were performed on Vanquish Neo nanoLC system coupled with an Orbitrap Eclipse mass spectrometer, a FAIMS Pro Interface, and an Easy Spray ESI source, all by Thermo Fisher Scientific. NanoLC separation utilized an Acclaim PepMap trap column (75 μm x 2 cm) combined with an EasySpray ES902 column (75 µm x 25 cm, 100) from Thermo Fisher Scientific. The injection volume for the peptide extrcts was 5 µL. Peptide separation was conducted with a mobile phase consisting of 0.1% (v/v) formic acid in water (solution A) and 0.1% (v/v) formic acid in 80% (v/v) acetonitrile (solution B), flowing at 300 nL/min, while the column temperature was held constant at 40°C. The column underwent initial conditioning with 3% solution B, followed by a gradual linear gradient up to 25% solution B over a 90 min period. Any remaining peptides adhering to the C18 resin were subsequently washed off with 95% solvent B for 8.5 min.

The mass spectrometer was operated in positive mode. The ion source temperature was set to 305°C, and ionized peptides were passed through the FAIMS Pro unit at -45, -55, and -65 V. Mass spectra were collected in MS1 mode with a resolution of 120,000, spanning the mass range of 375-1500 m/z. This was carried out using custom automatic gain control (AGC = 300) settings and automatic injection time. For MS2 data acquisition, the MS was operated in DDA mode with dynamic exclusion and a resolution of 30,000. The top precursors with charge states 2-7 were selected for MS2 analysis using a dynamic exclusion window of 45 sec. The following parameters were used for MS2 acquisition: dynamic maximum injection time, standard AGC target, isolation window of 1.6 m/z, 30% normalized collision energy, 30,000 resolution, and an intensity threshold of 2.0E4.

The Fragpipe 19.0 software package (version 19.0, MSFragger version 3.6, IonQuant version 1.8.9, Philosopher version 4.7.0) was used to analyze proteome data. The automatic centroiding, deisotoping, and deconvolution steps were also applied. Protein identification was performed using a FASTA database for *Bos taurus* proteins (Uniprot accession UP000009136) under the specified parameters, including:

- Parent mass error tolerance: 20 ppm
- Fragment mass error tolerance: 20 ppm
- Enzyme specificity: Semi-specific trypsin (cleavage at C-terminal of K/R with allowed 2 missed cleavages)

Additionally, carbamidomethylation (C) was set as an unvariable modification, while the following variable modifications were considered: oxidation (M) and acetylation (N-terminus). A non-specific cleavage on one terminus was permitted, with a maximum three variable post-translational modifications per peptide. The decoy-fusion approach was used to estimate false discovery rates (FDRs), and peptides with an FDR ≤ 1% and 1 unique peptide were regarded as confidently identified. Peptides considered for label-free quantification were selected using a precursor m/z tolerance of 10 ppm and a RT tolerance of 1 min.

For protein quantification, the two most abundant peptide signals were used to calculate raw protein peak areas. Quantitative protein data were exported in .csv format and further analyzed according to the methods described by Schulte *et al.*^69^.

### Acridine orange staining and FACS

Cells were stained with 4 ug/mL acridine orange (Invitrogen A3568) before FACS or analysis by flow cytometry.

### Bioanalyzer

RNA was analyzed on an Agilent Bioanalyzer using instructions from the manufacturer. Less than 5 ng of RNA was loaded on RNA pico chips (Agilent 5067-1513). The plots shown always come from samples that were treated identically and analyzed on the same bioanalyzer chip.

### RNA sequencing

Purified RNA (ranging from 100 to 2000 ng) from whole-cell and lysosomal samples had equal amounts of SIRV-Set 3 (Iso Mix E0/ERCC) (Lexogen 051.01) spike-ins added to them such that spike-ins would represent ∼0.01 to ∼0.2% of reads. They were then treated with DNAse I (Zymo E1010) according to Zymo’s instructions. RNA was then purified using Trizol LS and zymo-RNA clean and concentrators kits as above. Then, RNA was fragmented for 7 min at 94°C using NEB RNA fragmentation module (E6150S), before again purifying RNA using Trizol LS and zymo-RNA clean and concentrators kits as above. Then, to remove any 3′ cyclic phosphates from fragmentation, 30 uL of RNA was PNK treated with 50 U T4 PNK (Biosearch technologies P0503K) in a final volume of 60 uL prior to again purifying RNA using Trizol LS and zymo-RNA clean and concentrators kits as above. RNA was then quantified and checked for proper fragmentation by bioanalyzer as outlined above.

RNA sequencing libraries from these samples were prepared based on Xu *et al.*^38^ with some modifications. Briefly, reverse transcription was done using TGIRT-III from the Lambowitz lab using an annealed R2 RNA (rArArGrArUrCrGrGrArArGrArGrCrArCrArCrGrUrCrUrGrArArCrUrCrCrArGrUrCrArC/3S pC3/) / R2R DNA (GTGACTGGAGTTCAGACGTGTGCTCTTCCGATCTTN) mix. DNA was cleaned up using MinElute reaction cleanup kits (Qiagen 28204). Then cDNA was ligated to adenylated (NEB E2610S) R1R DNA (/5Phos/NNNNNNNNNNGATCGTCGGACTGTAGAACTCTGAACGTGTAG/3SpC3/) using Thermostable 5′ AppDNA/RNA ligase (NEB M0319S). DNA was cleaned up using MinElute reaction cleanup kit.

cDNA was amplified using Phusion (Thermo Fisher F531S) and the following primers: P5 barcode primer: AATGATACGGCGACCACCGAGATCTACAC-BARCODE- GTGACTGGAGTTCAGACGTGTGCTCTTCCGATCTT, P7 barcode primer: CAAGCAGAAGACGGCATACGAGAT-BARCODE- CTACACGTTCAGAGTTCTACAGTCCGACGATC. A cycle course of (6, 9, 12, 15, or 18 cycles) or (8, 10, 12, 14, or 16 cycles) was performed on half of the cDNA to determine the fewest amount of cycles necessary for library amplification (established by the PCR product being visible on an ethidium bromide stained agarose gel). Note, that here the primers used are different than those in the Lambowitz protocol. This is for two reasons. One is to use the non-truncated p5 sequence, and the other is to invert the reads such that the base biased sequence near the barcode is read second. If this is not done, sequencing quality is dramatically diminished, because NovaSeq is highly sensitive to base bias in the first 26 cycles.

PCR products were cleaned up to remove primers using 2 rounds of ampure xp bead (Beckman Coulter A63880) with a 1.4 x ratio of beads to PCR volume prior to submission for paired end 50nt sequencing on NovaSeq. As a result of the differences in primer design outlined above, sequencing was done with custom oligos. Read 1 primer was oGJR992: GTGACTGGAGTTCAGACGTGTGCTCTTCCGATCTT. Read 2 primer was oGJR993: CTACACGTTCAGAGTTCTACAGTCCGACGATC. i7 sequencing primer was oGJR994: GATCGTCGGACTGTAGAACTCTGAACGTGTAG. i5 sequencing primer was oGJR995: GATCGGAAGAGCACACGTCTGAACTCCAGTCAC.

Fastq files from multiple lanes were concatenated and them trimmed using fastp^70^ with the command:

fastp -i R1.fq.gz -I R2.fq.gz -o trimmed.R1.fq.gz -O trimmed. R2.fq.gz -U -- umi_loc=read2 --umi_len=10 -- adapter_sequence=GATCGTCGGACTGTAGAACTCTGAACGTGTAG -- adapter_sequence_r2=AAGATCGGAAGAGCACACGTCTGAACTCCAGTCAC

The rRNA “genome” was built with the following command:

STAR --runMode genomeGenerate --genomeSAindexNbases 6 --genomeDir STAR_rRNA --genomeFastaFiles 5S.fasta 5_8S.fasta 18S.fasta 28S.fasta --sjdbGTFfile rRNA.gtf --sjdbOverhang 39 --runThreadN 16. Fasta files are from NCBI and rRNA.gtf was created using biopython^71^ to create a gff file from a fasta and gffread^72^ to make a gtf from a gff file.

The tRNA “genome” was built with the following command:

STAR --runMode genomeGenerate --genomeSAindexNbases 5 --genomeDir STAR_tRNA --genomeFastaFiles CCA_hg38-tRNAs.fasta --sjdbGTFfile CCA_hg38- tRNAs.gtf --sjdbOverhang 39 --runThreadN 16. A tRNAs fasta of sequences was from the genomic tRNA database^73^ but had CCAs added to their 3′ using biopython such that mature tRNAs were not discarded for containing sequence that does not match the reference. GTF files were created as above.

The mitochondrial RNA “genome” was built with the following command: STAR --runMode genomeGenerate --genomeSAindexNbases 8 --genomeDir

STAR_mito --genomeFastaFiles mito.fasta --sjdbGTFfile mito.gtf --sjdbOverhang 39 -- runThreadN 16. The fasta and gff files are from NCBI reference NC_012920.1. GTF files were created as above.

The ncRNA “genome” was built with the following command:

STAR --runMode genomeGenerate --genomeSAindexNbases 13 –genomeDir STAR_ncRNA --genomeFastaFiles Min_Nc_RNA_Fasta_transcripts.fasta --sjdbGTFfile Min_Nc_RNA_Fasta_transcripts.gtf --sjdbOverhang 39 --runThreadN 16. Here, we wanted to map to abundant ncRNAs that may have homologs or pseudogenes throughout the genome. To do so we used biopython to first only look at RNAs not annotated as pseudogenes and then filtered for any RNAs with the biotype snRNA, snoRNA, or scaRNA as well as the RNAs RNY1, RNY3, RNY4, RN7SL1, RN7SL2, RN7SL3, VTRNA1, VTRNA2 and RPPH1. GTF files were created as above.

The “genome” for the whole transcriptome with the spike-in RNAs was built with the following command:

STAR --runMode genomeGenerate --genomeDir STAR_hg38_ERCC_SIRV -- genomeFastaFiles hg38_Gencode_v43_primary_ERCC_SIRV.fa --sjdbGTFfile hg38_Gencode_v43_primary_ERCC_SIRV.gtf --sjdbOverhang 39 --runThreadN 16. Fasta and GTF files for the primary GRCh38 version 43 human genome were downloaded from Gencode and concatenated with ERCC_SIRV fasta and GTF files downloaded from Lexogen. For the 5EU experiment fastas and GTFs with the sequences of nLUC and GFP were also included.

Reads were then mapped using STAR^74^ successively such that unmapped reads were then remapped to the next “genome” in the following order: rRNA, tRNA, mitochondrial RNA, ncRNA, and the whole transcriptome with the spike-in RNAs added. Then featurecounts^75^ was used with the following command for rRNA, mitochondrial RNAs, and the whole transcriptome with spike-in RNAs: featureCounts -p -d 15 -s 2 -T 16 -t exon -B -C. Because many RNAs are highly similar for tRNA and ncRNAs, reads were allowed to multimap so the following command was used: featureCounts -M --fraction -p -d 15 -s 2 -T 16 -t exon -B -C. In this instance a multimapping read was counted as 1/number of locations that it maps to. Using R, reads were then combined from all mappings and filtered for distinct geneids keeping the first instance such that redundant genes in multiple datasets were removed and the one first mapped to was retained. Then reads were normalized to their length /1000 to get read per kilobase (RPK) values.

For lysosomal enrichment experiments, ERCC and SIRV spike in values were filtered for either having 18/24 or 16/16 samples having 10+ reads. Then reads for each sample and the corresponding known spike-in concentrations from Lexogen were fit with a linear regression model. Then RPK values were divided by the slope value from the linear regression, and subsequently also multiplied by the values below to account for both differences in lysosomal capture, as quantified by LAMP2 western intensities, as well as normalizations for how much material was input into the IP versus taken for whole-cell and if RNA was diluted prior to spike-in addition. Normalization factors for the WT, KO, tKO lysosomal enrichment experiment were as follows. Whole cell samples: 14285, tKO lysoIP reps 1 & 2: 1025 & 775, KO lysoIP reps 1 & 2: 45 & 57, WT lysoIP reps 1 & 2: 47 & 46, tKO Torin lysoIP reps 1 & 2: 859 & 1000, KO Torin lysoIP reps 1 & 2: 295 & 459, and WT Torin lysoIP reps 1 & 2: 41 & 66. Normalization factors for the ATG7, FIP200, and LARP1 KO lysosomal enrichment experiment were as follows.

Whole cell samples: 1800, control KO LysoIP reps 1 & 2: 112 & 90, ATG7 KO LysoIP reps 1 & 2: 89 & 104, FIP200 KO LysoIP reps 1 & 2: 100 & 97, and LARP1 KO LysoIP reps 1 & 2: 83 & 93. To calculate lysosomal enrichment for each genotype, the average normalized lysosomal reads of the 2 biological replicates was divided by the average normalized whole cell reads of the 2 biological replicates. For the WT, KO, tKO lysosomal enrichment experiment the replicates used are from independently generated clonal cell lines. To define the role of ATG7 in lysosomal RNA localization, we calculated the average lysosomal enrichment in tKO cells that express ATG7 targeting guide RNAs and Cas9, and divide that by the average lysosomal enrichment of the 2 biological replicates in tKO cells that express control targeting guide RNAs and Cas9.

The same analysis was applied to FIP200 and LARP1.

For tRNA specific analysis, reads from tRNAs with identical sequences were combined and hydropathicity values were from (https://web.expasy.org/protscale/pscale/Hydropath.Doolittle.html)^42^.

### 5EU labeling and isolation of labeled RNA

150 mL of cells were incubated with 0.5 mM 5EU for 10 min shaking at 37°C, then spun at 300 x g for 3 min, and resuspended in 50 mL of media with 1 mM uridine (∼20 min until 5EU was removed). Cells were then spun 2 more times as above resuspending in 50 mL media with 1 mM uridine, and then 200 mL media with 1 mM uridine. Then cells were incubated at for 2 h post initial wash, with shaking at 37°C. At this point Torin1 (Selleck Chemicals S2827) treatment was started on half of the samples. Timepoints were collected by spinning down 15 mL cells at 300 x g for 3 min and resuspending in 4 mL of Tri reagent (Thermo Fisher Scientific AM9738) and freezing at -80°C.

Cells in tri-reagent were thawed and had 1 ng GFP and 5 ng of nLuc spike-in RNAs added to them. Spike-in RNAs were made by *in vitro* transcription as in Eisen *et al.*^21^. RNAs were then precipitated by adding 100 uL BCP for each 1 mL of tri reagent, mixing by shaking for 15 s, incubation at room temperature for 5 to 15 min, and then centrifugation at 12,000 x g for 15 min at 4°C. The aqueous phases (490-500 ul) were transferred to a fresh tube and 1 uL of 5 mg/mL linear acrylamide was added to each sample. 500 ul of isopropanol was then added per 1 mL of Tri reagent solution used for the sample. The samples were then mixed by vortexing and incubated at -20°C overnight. Samples were centrifuged at 12,000 x g for 8 min at 4°C, and the supernatant was removed without disturbing the pellet. Samples were washed with 1 mL cold 75% ethanol, then centrifuged at 12,000 x g for 5 min at 4°C. Ethanol was removed, and samples were centrifuged again briefly to remove any residual ethanol. Samples were air dried for 5 min and then dissolved in 100 ul water or 150ul water for the four untreated 28 h timepoints.

Biotinylation reactions were done as Eisen *et al.*^21^. Briefly, half of the isolated RNA from above was incubated in a 50 mM <u>HEPES</u>, pH 7.5, 4 mM disulfide <u>biotin</u> azide (Click Chemistry Tools), 2.5 mM CuSO_4_, 2.5 mM Tris(3-hydroxypropyltriazolylmethyl)amine (THPTA, Sigma-Aldrich), and 10 mM sodium ascorbate (Sigma-Aldrich) in a final volume of 180 uL for 1 h at room temperature protected from light. Reactions were stopped by adding EDTA to a final concertation of 5 mM, brought to 300 uL with water, and then had 300 uL phenol:chloroform:isoamyl alcohol (25:24:1,Sigma P2069-100ML) added. Samples were mixing by vortexing for 20 s, incubated at room temperature for 10 min, and centrifuged at 21,000 x g for 10 min at 4°C. The upper aqueous phase (280 uL) was transferred to a new tube, and 18 uL of 5 M NaCl, 2 uL 5 mg/mL linear acrylamide, and 900 uL ethanol were added, mixed, and incubated at -20°C overnight. Samples were then centrifuged at 21,000 x g for 10 min at 4°C and the supernatant was removed without disturbing the pellet. Samples were washed with 1 mL cold 75% ethanol, then centrifuged at 21,000 x g for 5 min at 4°C. Ethanol was removed, and samples were centrifuged again briefly to remove any residual ethanol. Samples were air dried for 5 min and then dissolved in 100 ul water.

RNA was fragmented for 7 min at 94°C using the NEB RNA fragmentation module (E6150S) in a final volume of 110 uL, and 10 uL of 3M pH 5.5 sodium acetate, 2 uL 5 mg/mL linear acrylamide, and 325 uL ethanol were added, mixed, and incubated at -20°C overnight. Samples were then centrifuged at 21,000 x g for 15 min at 4°C and supernatant was removed without disturbing the pellet. Samples were washed with 1 mL cold 70% ethanol, then centrifuged at 21,000 x g for 10 min at 4°C. Ethanol was removed, and samples were centrifuged again briefly to remove any residual ethanol. Samples were air dried for 5 min and then resuspended in 100ul 1X HSWB (10 mM Tris pH 7.5, 1 mM EDTA, 100 mM NaCl, 0.01% Tween-20).

Purification of biotinylated RNAs were done as Eisen *et al.*^21^. Dynabeads MyOne <u>Streptavidin</u> C1 beads (Invitrogen 65002) for each set of samples were combined and batch washed, with 200 μL of beads per reaction. Beads were washed twice with 1X B&W buffer (5 mM Tris-HCl, pH 7.5, 0.5 mM EDTA, 1 M NaCl and 0.005% Tween-20), twice with solution A (0.1 M NaOH, 50 mM NaCl), twice with solution B (0.1 M NaCl), and then twice with water, using for each wash a volume equal to that of the initial bead suspension. Following the last wash, beads were resuspended in an initial bead volume of 1X high salt wash buffer (HSWB, 10 mM Tris-HCl, pH 7.4, 1 mM EDTA, 0.1 M NaCl, 0.01% Tween-20) supplemented with 0.5 mg/mL yeast RNA (Thermo Fisher) and incubated at 23°C on a thermal mixer, shaking in a cycle of 15 s 1400 RPM on and 1 min 45 s off, for 30 min, again using a volume equal to that of the initial bead suspension. Beads were then washed three times with 1X HSWB per reaction and split for each reaction during the last wash. After the wash was removed, sample RNA previously resuspended in 100 μL 1X HSWB was added to blocked beads and incubated at 23°C on a thermal mixer, shaking in a cycle of 15 s 1400 RPM on and 1 min 45 s off, for 30 min. Beads were washed twice with 400 μL water at 50°C, incubating at 50°C for 2 min for each wash, and then twice with 400 μL 10X HSWB, incubating at 50°C for 2 min for each wash. RNA was eluted from beads by incubating with 180 μL 0.5 M tris(2-carboxyethyl)phosphine (TCEP, Sigma-Aldrich) at 50°C shaking on a thermal mixer in a cycle of 15 s 1400 RPM on and 1 min 45 s off, for 30 min. The initial eluate was collected, and beads were resuspended in 150 μL water and eluted again, combining the two eluates for each sample. RNA from the eluate was then mixed with 22 uL 5 M NaCl, 3 uL 5 mg/mL linear acrylamide and 1 mL ethanol and incubate at -20°C overnight. Samples were then centrifuged at 21,000 x g for 30 min at 4°C and supernatant was removed without disturbing the pellet. Samples were washed with 1 mL cold 75% ethanol, then centrifuged at 21,000 x g for 10 min at 4°C. Ethanol was removed, and samples were centrifuged again briefly to remove any residual ethanol. Samples were air dried for 5 min and then resuspended in 20 ul water.

Samples were then PNK treated as done above for lysosomal enrichment RNA sequencing, brought up to 350 uL with water, and mixed with 350 uL of phenol:chloroform:isoamyl alcohol (25:24:1,Sigma P2069-100ML). Samples were mixing by vortexing for 20 s, incubated at room temperature for 10 min, and centrifuged at 21,000 x g for 10 min at 4°C. The upper aqueous phase (340 uL) was transferred to a new tube, and 340ul chloroform was mixed in before incubating at room temperature for 10 min and centrifugation at 21,000 x g for 10 min at 4°C. The upper aqueous phase (280 uL) was transferred to a new tube, and 18 uL of 5M NaCl, 2 uL 15 mg/mL glycoblue, and 900 uL ethanol were added, mixed, and incubated at -20°C overnight.

Samples were then centrifuged at 21,000 x g for 30 min at 4°C and supernatant was removed without disturbing the pellet. Samples were washed with 1 mL cold 75% ethanol, then centrifuged at 21,000 x g for 10 min at 4°C. Ethanol was removed, and samples were centrifuged again briefly to remove any residual ethanol. Samples were air dried for 5 min and then dissolved in 20ul water. 50 ng of RNA, or 10ul for mock IP samples, were then input into the TGIRT RNA sequencing protocol outlined above.

### Half-life calculation

Mapping and analysis of RNA sequencing was done as above, except that samples were not normalized by length, and samples were normalized to a nLuc spike-in RNA by dividing by the counts for all other genes by the counts for nLuc in each sample. Then, for each gene the reads at each time-point were normalize to timepoint 0 such that all RNAs started at 1.

The data was then fit in R using nlsLM to the following equation for each replicate: y = log_2_(Ce^-kx^) + z. y is the log_2_ number of normalized counts at time x. z is a background metric that is the max value of any of the 4 mock IP samples or 0.000005 divided by the spike-normalized reads at timepoint 0(pre-normalization to nLuc). When fitting for each gene, C and k were bound from 0 to infinity, with starting values of 2^max(y)^ and 1 respectively. For Torin1 treated samples C was fixed as the C value of the untreated sample since C value represents an initial value that should be independent of the treatment. The half-life was calculated as ln(2)/k with a maximum of 50 h. Only half-lives where the Pr(>|t|) <.05 for k and C where recorded. A synthesis rate S was calculated as C*k. Downstream analyses are the average of independently calculated half-lives for the 2 biological replicates.

### Northern Blotting

For each sample, RNA from equal proportions of whole cell and lysosomal immunoprecipitates (2–10 μg RNA) was mixed with an equal volume of 2x Loading Dye (8 M urea, 25 mM EDTA, 0.025% xylene cyanol, 0.025% bromophenol blue). Samples were heated to 80-95°C in a heat block for 5 to 10 min and spun down. Samples were resolved on a denaturing 6% polyacrylamide gel (UreaGel-SequaGel-System, National Diagnostics) run at 5 W for 15 min and 25 W for 70 min. The gel was then transferred onto a Hybond-NX membrane (RPN303T GE Healthcare) in 0.5X TBE buffer using a semi-dry transfer apparatus (Bio-Rad) (3 mA/cm^2^ of membrane) for 35 min. The RNA was then UV cross-linked to the membrane with 254 nm light and incubated with 42°C Ultra-hyb-oligo solution (Ambion) for at least 15 min at 42°C. DNA probes were labeled by incubation 1 uL T4 PNK (NEB M0201L), 1 uL 10uM oligo, 1 uL 10X T4 PNK buffer (NEB M0201L), 6 uL water, and 1 uL ^32^P -ATP (Revvity BLU035C001MC) at 37°C for at least 1 h. 2 different probes were used since the probe used for truncation mutants would have had mismatches to several of the point mutant SRP RNAs. The truncation mutant blot used oGJR1005: tcctccagcctcagcctcccgagtagct. The point mutant blot used oGJR1036: aagcgatcctccagcctcagcctcccga. 50 uL of water was added to the labeling reaction and probe was purified from unicorporated label with Micro Bio-Spin P30 Columns (Biorad 732-6250) by following the manufacturer’s instructions. Column-purified probe was added directly to the blot in Ultra-hyb-oligo solution and allowed to hybridize at 42°C overnight. The blot was washed with ∼20 mL of wash buffer (0.5% SDS and 2X SSC) twice for 5 min, and once for 30 min all at 42°C. The blot was put in a Kapak heat-seal bag and placed in a cassette with a blanked phosphorimager plate to expose. Results were analyzed on a Typhoon FLA 7000 phosporimager, using ImageQuant TL (v8.1.0.0) for quantification of band intensities and ImageJ (1.52q) for generation of smoothed line densitometry profiles.

### Transient transfections

The day prior to transfection, RNASET2 PLD3 RNASE4 tKO HEK293T cells were resuspended in fresh media at ∼1.5 million cells/mL in freestyle media without serum. The next day, cells were transfected with 1 ug of DNA per 1 mL of culture media and a 3:1 ratio of 1 mg/mL PEI STAR (Bio-techne 7854) to total DNA. Only 10% of DNA was the plasmid of interest, and the rest was salmon sperm DNA (Invitrogen 15632011) that had been heated to 95°C for 10 min and then incubated on ice for 10 min prior to mixing with other components. For a 20 mL culture, 20 ug DNA was diluted and mixed in 1 mL of serum-free Opti-MEM media prior to addition of 60 uL of 1 mg/mL PEI STAR. The tube was inverted several times to mix and incubated at room temperature for 10 min prior to addition to the flask while swirling. The next day, media was added to a final volume of 50 mL, and cells were subsequently cultured normally until collection for lysoIP.

#### RT-qPCR

For each sample, 5 uL of total RNA was reverse-transcribed using super-script III (Invitrogen 18080044) with random-hexamer priming, followed by qPCR using SYBR Green PCR Master Mix (Applied Biosystems 4309155) on a QuantStudio 6 system (Applied Biosystems). Reporter transcripts were quantified using the ΔΔCT method with 18S rRNA levels as internal controls. Primers for GFP were oGJR129:GCACAAGCTGGAGTACAACTA and oGJR130:TGTTGTGGCGGATCTTGAA. Primers for 18S rRNA were oGJR478:agacaaatcgctccaccaac and oGJR479:cctgcggcttaatttgactc.

## Acknowledgments

We thank members of the Sabatini and Bartel laboratories for discussions and feedback, especially Arash Latifkar, Julian Roessler, Anna Traunbauer, Brad Wierbowski, and Kehui Xiang. We also thank L. Bryan Ray, Jill M. Ray, and Frank Solomon for feedback on the manuscript. We thank Alan Lambowitz’s lab for TGIRT enzyme. We thank the Whitehead Genome Technology Core for sequencing, and the Whitehead Quantitative Proteomics Core for proteomics. This work was supported by grants from the NIH (T32 GM007287 and F31 GM129905 to G.J.R.; R01 CA103866, R01 CA129105, and R01 AI047389 to D.M.S.), the Department of Defense (W81XWH-21-1-0260 and TS200035 to D.M.S.), the Lustgarten Foundation to D.M.S., the Leo Foundation to D.M.S., the Institute of Organic Chemistry and Biochemistry of the Czech Academy of Sciences to D.M.S., Pershing Square Philanthropies to D.M.S., and the MIT School of Science Fellowship in Cancer Research from the Koch institute for Integrative Cancer Research at MIT to G.J.R. D.H.L was an HHMI fellow of the Damon Runyon Cancer Research Foundation (DRG-2345-18). D.P.B. is an investigator of the Howard Hughes Medical Institute.

## Author contributions

Conceptualization: G.J.R., D.M.S., D.P.B.; Methodology: G.J.R.; Validation: G.J.R., E.N.; Formal Analysis: G.J.R.; Investigation: G.J.R., D.H.L., H.R.K.; Resources: G.J.R., E.N.; Writing – Original Draft: G.J.R.; Writing – Review & Editing: G.J.R., E.N., D.H.L., H.R.K., D.M.S., D.P.B.; Supervision: D.M.S., D.P.B.; Funding Acquisition: G.J.R., D.M.S., D.P.B.

## Materials & Correspondence

Correspondence and material requests can be made to G.J.R., D.M.S., and D.P.B.

## Data Availability

Sequencing data was deposited in Gene Expression Omnibus (accession number GSE296736), proteomics data are available via ProteomeXchange with identifier PXD063773, and plasmids will be submitted to Addgene.

---

**Extended Data Fig. 1:**
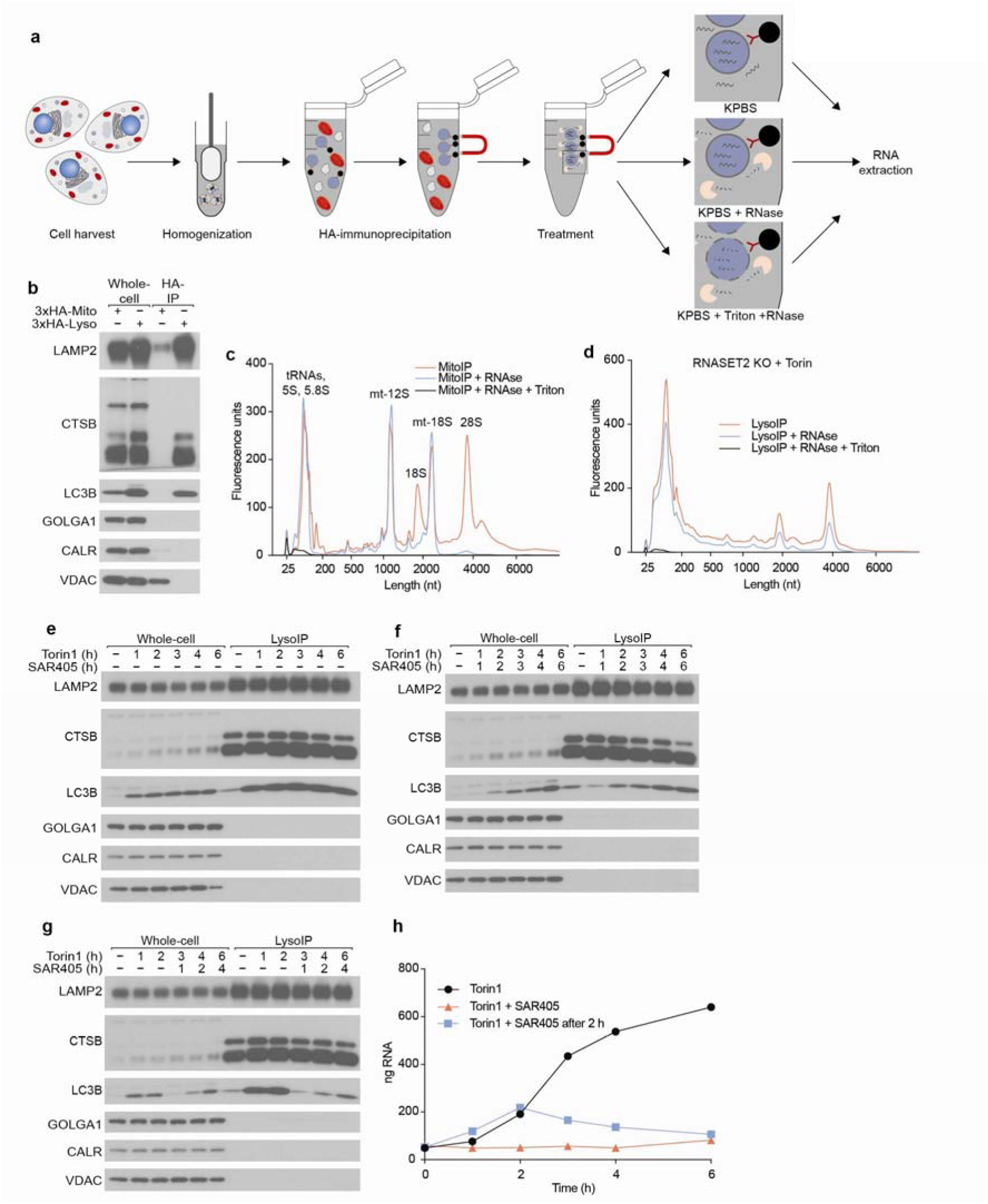
Purification strategy to obtain luminal organellar RNAs and the dependence of lysosomal RNA accumulation in RNASET2-KO cells on Torin1 and SAR405, related to Fig. 1. **a**, Schematic of organellar RNA purification. HEK293T cells were dounce-homogenized to disrupt their plasma membrane while keeping organelles intact, followed by an HA-immunoprecipitation to isolate organelle of interest. Then, purified organelles were subjected to either no treatment, treatment with RNase I alone, or treatment with RNase I in the presence of the detergent Triton X-100, followed by RNA extraction with TRIzol LS. **b**, Lysosomal and mitochondrial purification efficiencies. Organellar capture efficiency and purity were assessed by immunoblotting for protein markers of various subcellular compartments in whole-cell lysates, purified mitochondria, or purified lysosomes. Equal numbers of HEK293T cells stably expressing 3x-HA-EGFP-OMP25 (3xHA-Mito) or TMEM192-3x-HA (3xHA-Lyso) were subjected to the organellar IP protocol. 3xHA-Lyso cells were RNASET2-KO cells treated with 250 nM Torin1 for 4 h. **c**, RNase membrane protection assay on MitoIP purified mitochondria. Mitochondrial immunoprecipitates from (**b**) were subjected to either no treatment, treatment with RNase I alone, or treatment with RNase I in the presence of the detergent Triton X-100. Then, RNA was purified and resolved on a bioanalyzer. The peak at 25 nt is a marker. Other well-established peaks are tRNAs and cytosolic and mitochondrial (mt) rRNAs, as marked above each peak. **d**, RNase membrane protection assay on LysoIP purified lysosomes. Lysosomal immunoprecipitates from (**b**) were subjected to either no treatment, treatment with RNase I alone, or treatment with RNase I in the presence of the detergent Triton X-100. Then, RNA was purified and resolved on a bioanalyzer. The peak at 25 nt is a marker. **e**, Lysosomal purification efficiencies during a Torin1 time course. Lysosomal capture efficiency and purity were assessed by immunoblotting for protein markers of various subcellular compartments in either whole-cell lysates or purified lysosomes. Equal numbers of RNASET2-KO HEK293T cells stably expressing TMEM192-3x-HA were subjected to the LysoIP protocol. Where indicated, cells were treated with Torin1 at 250 nM. **f**, Lysosomal purification efficiencies during a Torin1 & SAR405 time course. This panel is as in (**e**), except cells were also treated with SAR405 at 3 uM. **g**, Lysosomal purification efficiencies during a Torin1 and delayed SAR405 time course. This panel is as in (**f**) except SAR405 treatment was delayed until 2 h after Torin1 addition. **h**, RNA quantifications across time courses above. Lysosomal RNAs from (**e–g**) were quantified by bioanalyzer.

**Extended Data Fig. 2:**
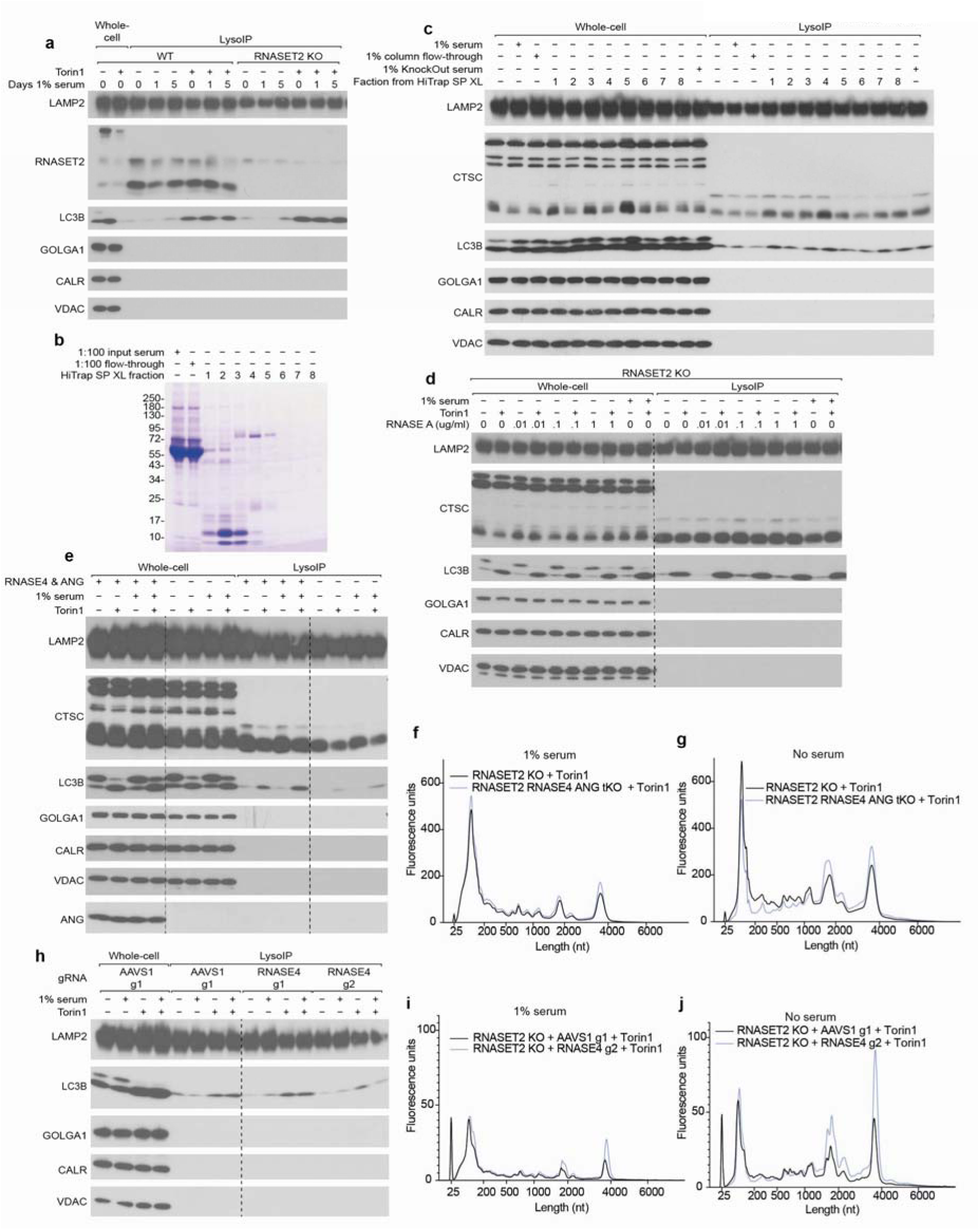
Endogenous and exogenous RNase A contribute to lysosomal RNA degradation, related to Fig. 3. **a**, Lysosomal purification efficiencies across serum and Torin1 treatments. Lysosomal capture efficiency and purity were assessed by immunoblotting for protein markers of various subcellular compartments in whole-cell lysates, or purified lysosomes. Equal numbers of HEK293T cells stably expressing TMEM192-3x-HA were subjected to the LysoIP protocol. Where indicated, cells were treated with 250 nM Torin1 for 5 h. **b**, Coomassie blue analysis of serum fractionated by cation exchange. Bovine serum was run through a HiTrap SP XL column, eluted using a gradient of NaCl, and fractions were collected and desalted prior to analysis by gel electrophoresis. **c**, Lysosomal purification efficiencies across fractionated serum treatments. Lysosomal capture efficiency and purity were assessed by immunoblotting for protein markers of various subcellular compartments in whole-cell lysates, or purified lysosomes. Equal numbers of RNASET2 PLD3 double-KO HEK293T cells stably expressing TMEM192-3x-HA grown in serum-free media had serum or fractionated serum added to the culture overnight prior to a LysoIP the next day. **d**, Lysosomal purification efficiencies across supplemental RNAse treatments. Lysosomal capture efficiency and purity were assessed by immunoblotting for protein markers of various subcellular compartments in whole-cell lysates, or purified lysosomes. Equal numbers of RNASET2-KO HEK293T cells stably expressing TMEM192-3x-HA grown in serum-free media had serum or RNase A added to the culture overnight prior to LysoIP the next day. Where indicated, cells were treated with 250 nM Torin1 for 5 h. **e**, Lysosomal purification efficiencies across the indicated genotypes and treatments. Lysosomal capture efficiency and purity were assessed by immunoblotting for protein markers of various subcellular compartments in whole-cell lysates, or purified lysosomes. Equal numbers of RNASET2-KO or RNASET2, RNASE4, ANG triple KO HEK293T cells stably expressing TMEM192-3x-HA, were subjected to the LysoIP protocol. Cells were grown in serum-free media and if indicated had serum added to the culture overnight prior to LysoIP the next day. Where indicated, cells were treated with 250 nM Torin1 for 6 h. **f**, Effect of RNASE4 and ANG on lysosomal RNA degradation in the presence of serum. Lysosomal RNAs from equal proportions of immunoprecipitates from cells cultured with or without RNASE4 and ANG (**e**) were purified and resolved on a bioanalyzer. The peak at 25 nt is a marker. Cells were grown in the presence of 1% serum overnight prior to LysoIP. Torin1 treatment was 250 nM for 6 h. **g**, Effect of RNASE4 and ANG on lysosomal RNA degradation in the absence of serum. Otherwise, this panel is as in (**f**). **h**, Lysosomal purification efficiencies across the indicated genotypes and treatments. Lysosomal capture efficiency and purity were assessed by immunoblotting for protein markers of various subcellular compartments in whole-cell lysates, or purified lysosomes. Equal numbers of RNASET2-KO HEK293T cells stably expressing TMEM192-3x-HA, Cas9, and the indicated guide RNA, were subjected to the LysoIP protocol. Cells were grown in serum-free media and if indicated had serum added to the culture overnight prior to LysoIP the next day. Where indicated, cells were treated with 250 nM Torin1 for 6 h. **i**, Effect of RNASE4 on lysosomal RNA degradation in the presence of serum. Lysosomal RNAs from equal proportions of immunoprecipitates from cells cultured with or without RNASE4 (**h**) were purified and resolved on a bioanalyzer. The peak at 25 nt is a marker. Cells were grown in the presence of 1% serum overnight prior to LysoIP. Torin1 treatment was 250 nM for 6 h. **j**, Effect of RNASE4 on lysosomal RNA degradation in the absence of serum. This panel is as in (**i**) except cells were grown in the absence of serum.

**Extended Data Fig. 3:**
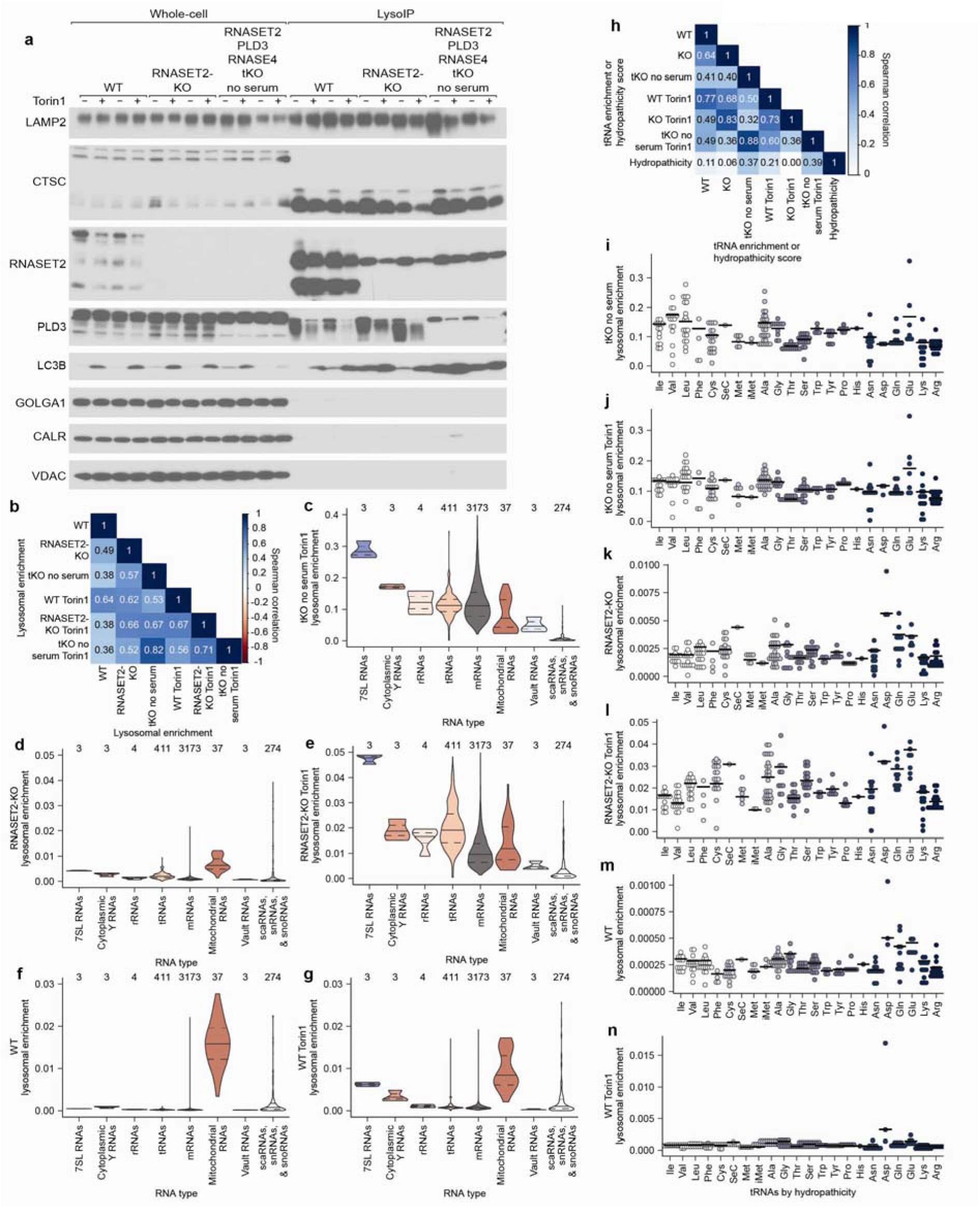
Specific RNAs are targeted for lysosomal degradation, related to Fig. 4. **a**, Lysosomal purification efficiencies across the indicated genotypes and treatments. Lysosomal capture efficiency and purity were assessed by immunoblotting for protein markers of various subcellular compartments in whole-cell lysates or purified lysosomes. Equal numbers of HEK293T cells stably expressing TMEM192-3x-HA were subjected to the LysoIP protocol. Where indicated, cells were treated with 250 nM Torin1 for 5 h. **b**, Spearman correlations between lysosomal enrichments. **c–g**, Lysosomal enrichments for different RNA types across the indicated genotypes and treatments. Where indicated, cells were treated with 250 nM Torin1 for 5 h. The number above each violin plot indicates how many unique RNAs were considered in that plot. **h**, Spearman correlations between lysosomal tRNA enrichments and amino acid hydropathicity^42^. **i–n**, Lysosomal enrichment of tRNAs across the indicated genotypes and treatments. tRNAs are displayed in order of decreasing hydropathicity^42^ of the amino acid carried by the tRNA. Each dot indicates the lysosomal enrichment for a unique tRNA sequence, and the bar indicates the overall enrichment for tRNAs that carry that amino acid. Where indicated, cells were treated with 250 nM Torin1 for 5 h.

**Extended Data Fig. 4:**
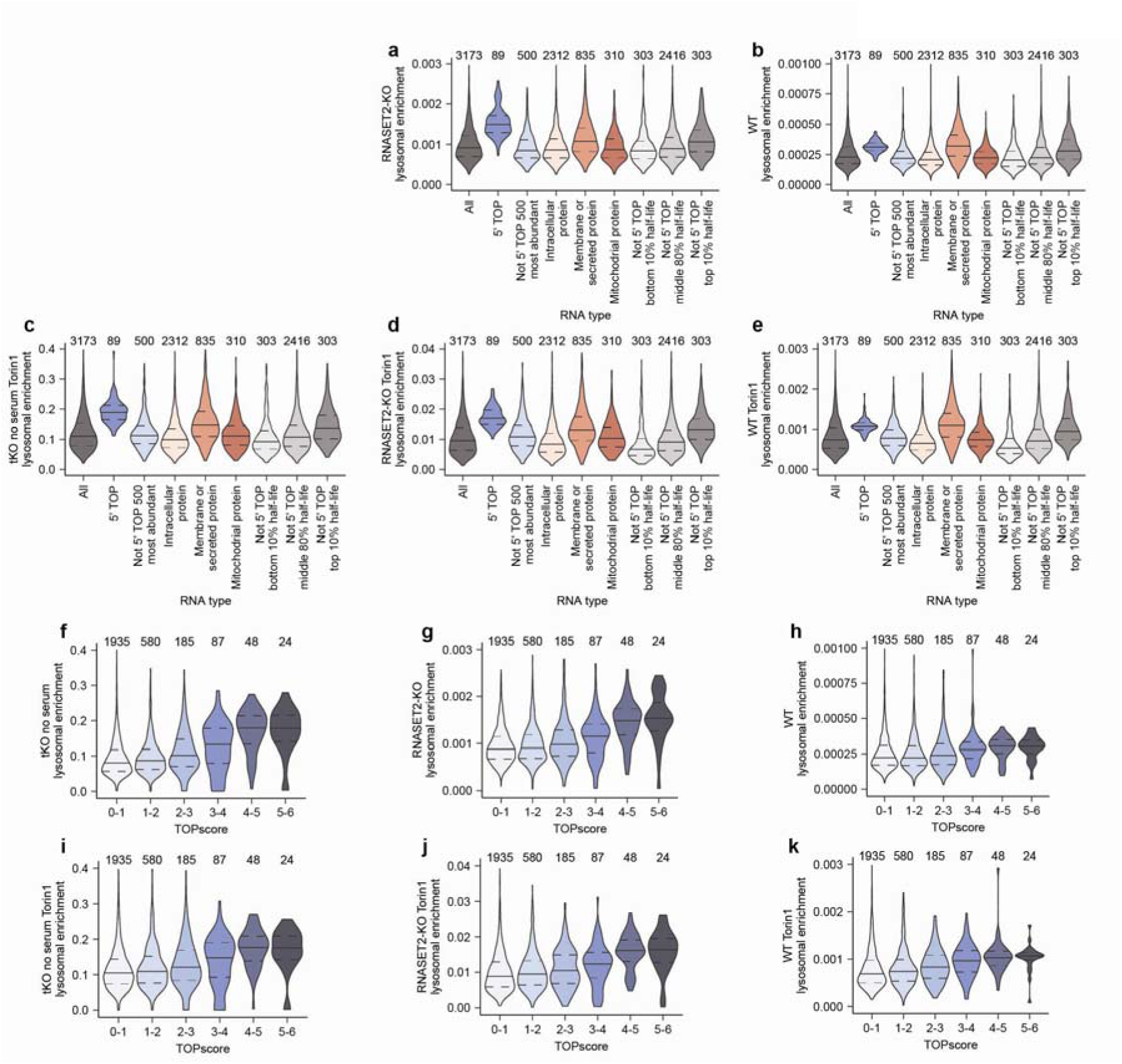
Specific mRNAs are targeted for lysosomal degradation, related to Fig. 4. **a– e**, Lysosomal enrichments for different mRNA types across the indicated genotypes and treatments. Protein localization data is from the Human Protein Atlas^39^, mitochondrial proteins were defined by Mitocarta3^40^, and half-life data is from Agarwal and Kelly^41^. The number above each violin plot indicates how many unique mRNAs were considered in that plot. Where indicated, cells were treated with 250 nM Torin1 for 5 h. **f–k**, Lysosomal enrichment binned by TOPscore^45^ across the indicated genotypes and treatments. Where indicated, cells were treated with 250 nM Torin1 for 5 h. The number above each violin plot indicates how many unique RNAs were considered in that plot.

**Extended Data Fig. 5:**
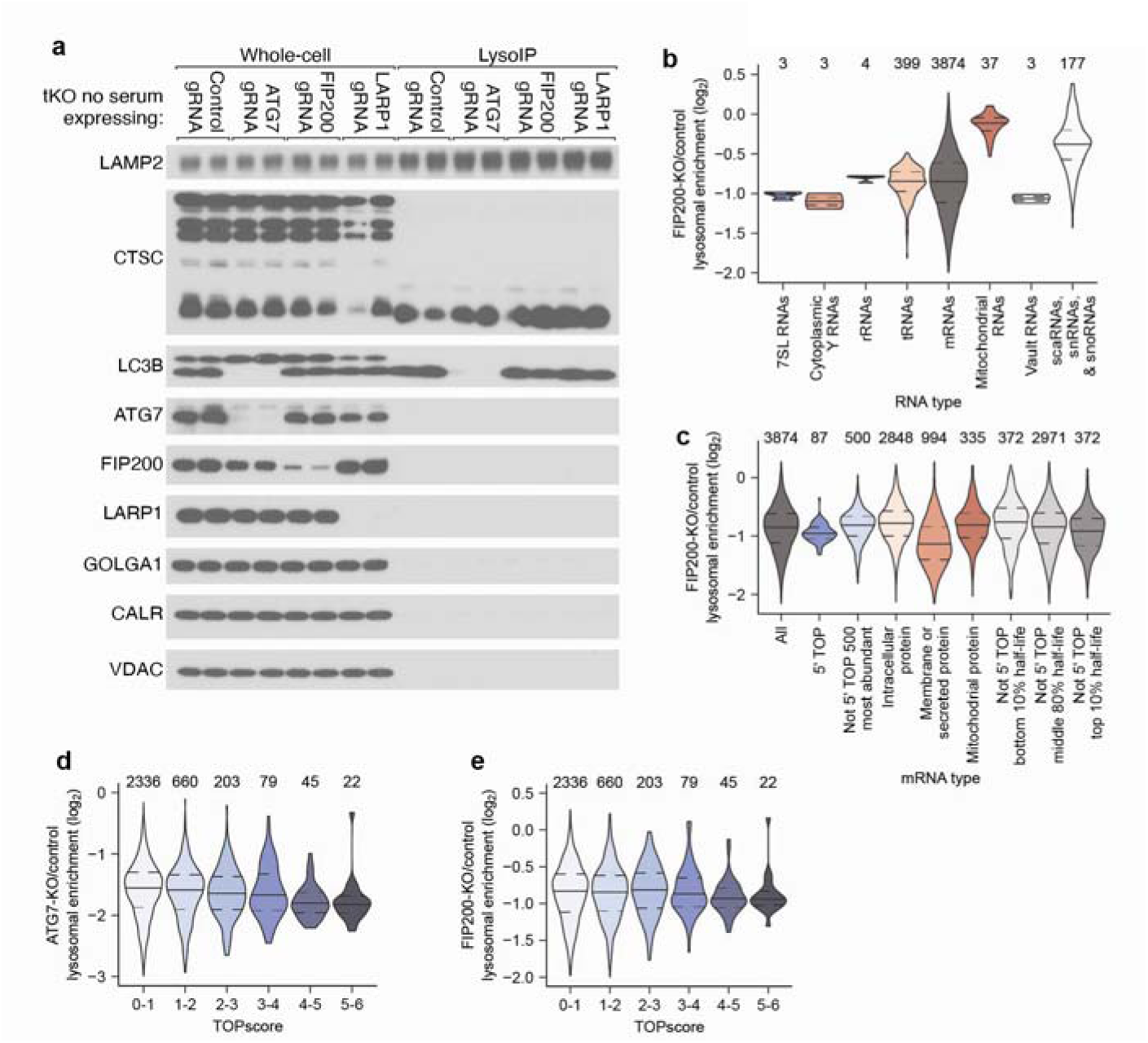
Autophagy targets specific RNAs for lysosomal degradation, related to Fig. 4. **a**, Lysosomal purification efficiencies across the indicated genotypes. Lysosomal capture efficiency and purity were assessed by immunoblotting for protein markers of various subcellular compartments in whole-cell lysates or purified lysosomes. Equal numbers of RNASET2, PLD3, RNASE4 tKO HEK293T cells stably expressing TMEM192-3x-HA, Cas9, and the indicated guide RNA, were subjected to the LysoIP protocol. **b**, Effect of FIP200 KO on lysosomal enrichment of different RNA types. The number above each violin plot indicates how many unique RNAs were considered in that plot. **c**, Effect of FIP200 KO on lysosomal enrichment of different mRNA types. Protein localization data is from the Human Protein Atlas^39^, mitochondrial proteins were defined by Mitocarta3^40^, and half-life data is from Agarwal and Kelly^41^. The number above each violin plot indicates how many unique mRNAs were considered in that plot. **d–e**, Effect of ATG7-KO (**d**) or FIP200-KO (**e**) on lysosomal enrichment as a function of the TOPscore^45^. The number above each violin plot indicates how many unique RNAs were considered in that plot.

**Extended Data Fig. 6:**
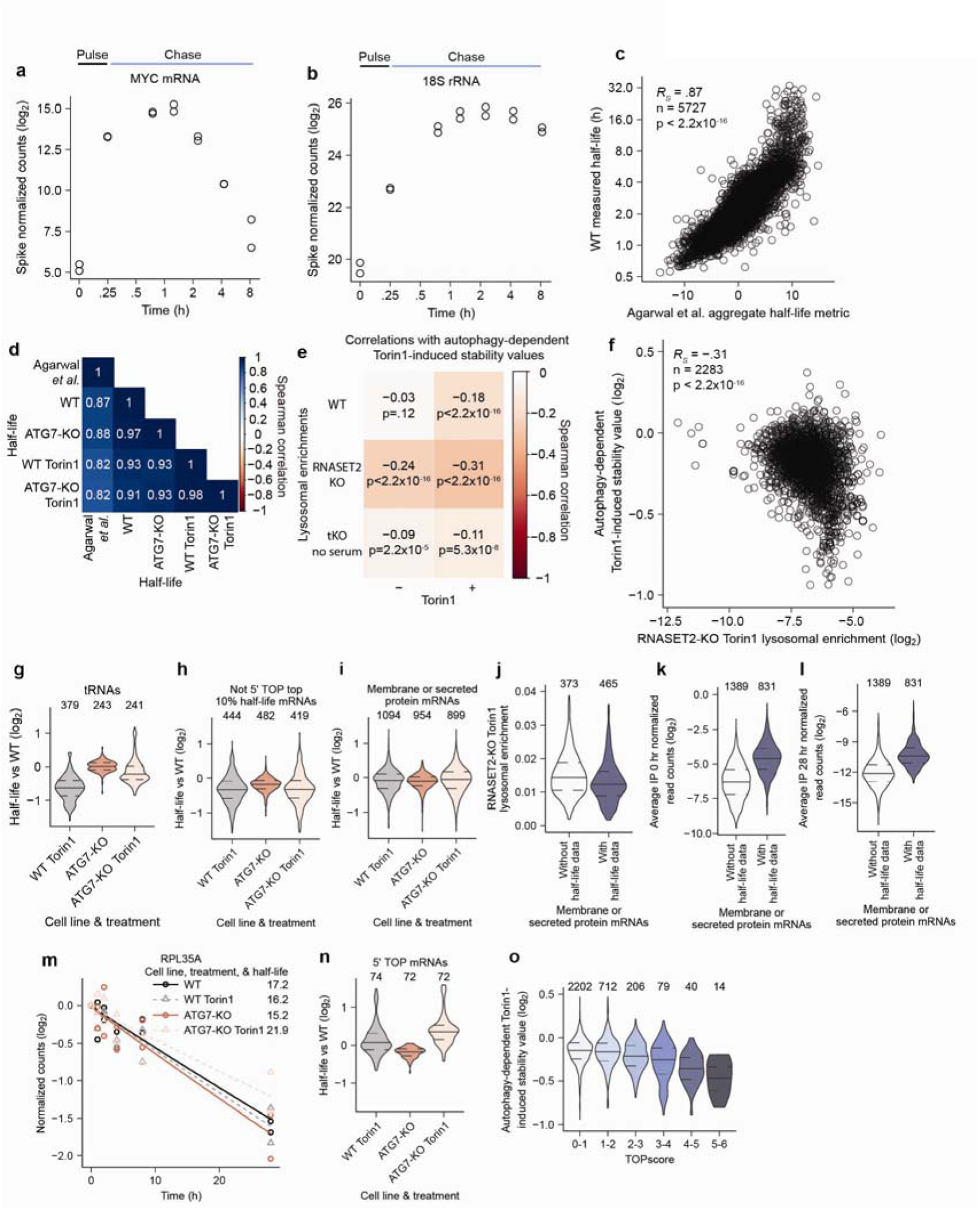
Establishing 5EU pulse-chase methods for RNA half-life measurements and the effects of autophagy and Torin1 on RNA stability, related to Fig. 5. **a**, Continued incorporation of 5EU into MYC mRNA transcripts in a conventional pulse-chase approach. **b**, Continued incorporation of 5EU into 18S rRNA transcripts in a conventional pulse-chase approach. **c**, Relationship between half-lives measured in wild-type cells and aggregate half-life metrics^41^. **d**, Spearman correlations between measured half-lives and aggregate half-life metrics^41^. **e**, Spearman correlations between autophagy-dependent Torin1-induced stability values and lysosomal enrichments. **f**, Relationship between autophagy-dependent Torin1-induced stability values and lysosomal enrichment in RNASET2-KO cells treated with Torin1. **g**, Effects of autophagy and Torin1 on tRNA stabilities. The number above each violin plot indicates how many unique RNAs were considered in that plot. **h**, Effects of autophagy and Torin1 on stabilities of stable mRNAs. This panel is as in (**g**) except RNAs analyzed are stable mRNAs determined by Agarwal and Kelly^41^. **i**, Effects of autophagy and Torin1 on stabilities of mRNAs encoding membrane and secretory proteins. This panel is as in (**g**) except mRNAs analyzed are those of membrane and secreted proteins, as annotated by the Human Protein Atlas^39^. **j**, Distributions of lysosomal enrichments in RNASET2-KO cells treated with Torin1 for mRNAs encoding lysosomally enriched membrane and secretory proteins, distinguishing between those that either do or do not have half-life measurements. The number above each violin plot indicates how many unique RNAs were considered in that plot. **k–l**, Fewer read counts for mRNAs encoding membrane and secretory proteins throughout the 5EU pulse-chase experiments. The number above each violin plot indicates how many unique RNAs were considered in that plot. **m**, Representative plot of effects of autophagy and Torin1 on the stability of a 5′ TOP mRNA. For each of the indicated genotypes and treatments, the lines represent decay equations determined by the average of decay rates and the average of an initial constant value established from biological replicates. **n**, Effects of autophagy and Torin1 on 5′ TOP mRNA stabilities. This panel is as in (**g**) except the mRNAs analyzed are 5′ TOP mRNAs. **o**, Autophagy-dependent Torin1-induced stability values binned by TOPscore^45^. The number above each violin plot indicates how many unique RNAs were considered in that plot.

**Supplementary Table 1 (separate file).** Results of the acridine orange CRISPR screen, related to Fig. 2.

**Supplementary Table 2. (separate file).** Proteomics of fractionated serum, related to Fig. 3.

**Supplementary Table 3. (separate file).** Lysosomal RNA sequencing, related to Fig. 4, and Extended Data Fig. 3, and 4.

**Supplementary Table 4. (separate file).** Lysosomal RNA sequencing, related to Fig. 4 and 7, and Extended Data Fig. 5.

**Supplementary Table 5. (separate file).** RNA stability measurements in WT and ATG7 KO cells, related to Fig. 5 and Extended Data Fig. 6.

